# The visual system of the longest-living vertebrate, the Greenland shark

**DOI:** 10.1101/2025.04.10.648203

**Authors:** Lily G. Fogg, Emily Tom, Maxime Policarpo, William Cho, Fangyuan Gao, Steven F. Grieco, Doreen Hii, Aaron E. Fawcett, Nicolas Boileau, Amalie Bech-Poulsen, Kirstine F. Steffensen, Cherlyn J Ng, Peter G. Bushnell, John Fleng Steffensen, Richard Brill, Walter Salzburger, Dorota Skowronska-Krawczyk

## Abstract

The Greenland shark (*Somniosus microcephalus*) is the longest-living vertebrate and inhabits the extremely dim and cold waters of the Arctic deep sea. This has led to speculations that it may have lost functional vision. Here, we present genomic, transcriptomic, histological and functional evidence that the Greenland shark retains an intact visual system well-adapted for life in dim light. Histology and *in vitro* opsin expression revealed visual adaptations typical of deep-sea species, including densely packed, elongated rods and a short-wavelength shift in rod visual pigment sensitivity. RNAscope confirmed the presence of essential visual cell types, such as rods, Müller glia, and bipolar, amacrine, and ganglion cells. Moreover, despite being centuries old, the examined specimens showed no signs of retinal degeneration. Using whole genome and retinal RNA-sequencing, we further show that dim-light (rod-based) vision genes are intact and robustly expressed, while many bright-light (cone-based) vision genes have become pseudogenized and/or are no longer expressed. Finally, our data suggest that efficient DNA repair mechanisms may contribute to the long-term preservation of retinal function over centuries in the Greenland shark.

## Introduction

The Greenland shark (*Somniosus microcephalus*; Fig. 1a) is the longest-living vertebrate on Earth, with an estimated lifespan of up to 400 years ^1^. This deep-sea shark inhabits regions from the temperate North Atlantic to the frigid waters of the Arctic Ocean, enduring temperatures as low as -1.1°C and depths approaching 3000 meters ^2-4^. In the waters surrounding Greenland, their eyes are frequently parasitized by copepods (*Ommatokoita elongata*), which are thought to obscure vision by attaching to the cornea ^5, 6^. The unique combination of extreme longevity, persistent low temperatures, high-pressure conditions, and parasitised eyes presents unparalleled challenges for its visual system – raising fundamental questions about the nature and function of the visual sensory system in this enigmatic species.

**Fig. 1.**
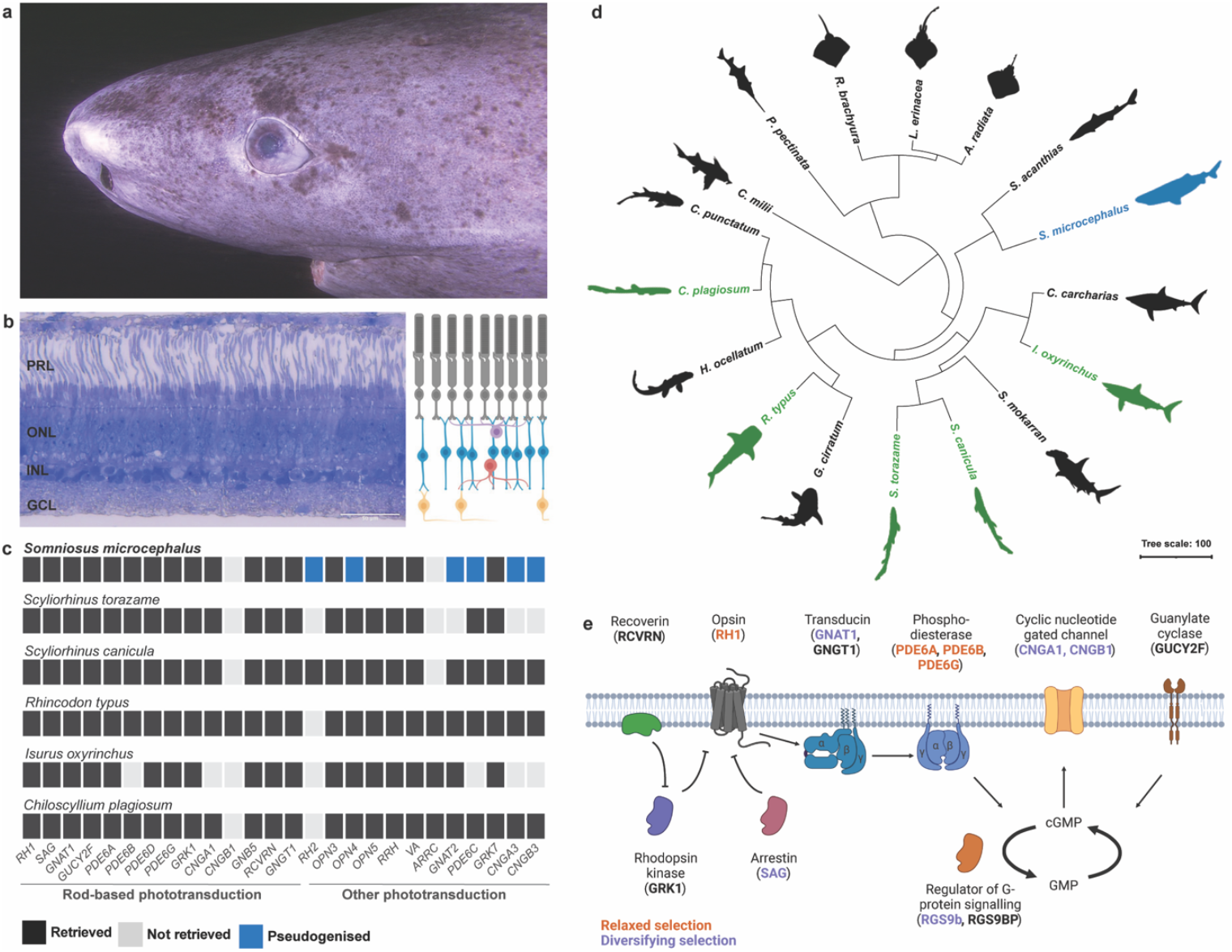
Visual system of the Greenland shark. **a**. Representative photograph of the head and eye of a Greenland shark, *Somniosus microcephalus*. **b**. Transverse section through the retina of the Greenland shark, alongside schematic illustration of the retinal layers. ***Scale bar:*** 50 µm. **c**. Tile plot summarising phototransduction gene mining. Note that most sharks (including the Greenland shark) lack a copy of *cngb1*. **d**. Species tree showing phylogenetic placement of the Greenland shark (blue), and five comparison species (green; *Scyliorhinus canicula, S. torazame, Isurus oxyrinchus, Rhincodon typus*, and *Chiloscyllium plagiosum*) used in this study within the class, Chondrichthyes. **e**. Schematic of the components of the rod-based phototransduction cascade found in the genome of the Greenland shark, with gene names coloured by evidence for relaxed (orange) or diversifying (purple) selection. ***Abbreviations***: PRL, photoreceptor layer; RPE, retinal pigment epithelium; ONL, outer nuclear layer; INL, inner nuclear layer; GCL, ganglion cell layer. Photograph in panel a was provided with permission for use by Ghislain Bardout from the Under the Pole expedition.

Vertebrate visual systems have evolved across diverse photic environments, from bright terrestrial habitats to the perpetual darkness of caves and the deep sea. In most vertebrate species, vision relies on two types of retinal photoreceptors: rods, optimized for low-light (scotopic) vision, and cones, specialized for bright-light (photopic) vision ^7^. Rods are highly sensitive, with densely packed chromatophore-bound visual pigments (opsins) and an efficient phototransduction cascade, while cones provide saturation-resistant, broad-spectrum photic sensitivity ^8^. The relative abundance of the photoreceptors and the spectral tuning of their opsins are primarily shaped by ecological pressures ^9-13^. While the rod opsin (RH1) is tuned to detect a narrow spectrum of blue-green light (460 – 530 nm), multiple cone opsins (SWS1, SWS2, RH2, and LWS) span a wide sensitivity range from ultraviolet to red light (350 – 600 nm) ^14^. Deep-sea and nocturnal species often exhibit rod-dominated retinas, sometimes to the complete exclusion of cones, as in certain nocturnal reptiles ^15^ and deep-sea teleost fishes ^16^.

In the most extreme conditions, vision can be entirely lost. Cavefishes, for instance, have evolved in complete darkness, leading to retinal degeneration, loss of phototransduction gene expression, and widespread pseudogenization of vision-related genes ^17-19^. Given the Greenland shark’s exceptionally dim and potentially obstructed visual environment, it has been speculated that it, too, may have lost its ability to see. However, behavioural observations suggest that these sharks may still rely on sight ^20^, and their optic tectum – a brain region that processes visual information – is comparable in size to that of other visually capable elasmobranchs ^21^. Furthermore, the Greenland shark has a tapetum lucidum, a specialized reflective layer behind the retina that enhances photon capture in low-light conditions ^22^.

In this study, to determine whether the Greenland shark possesses a lifelong functional visual system, we conducted a comprehensive integrative analysis incorporating genomics, transcriptomics, RNAscope, ultramicrotomy, chromatin staining, *in vitro* opsin regeneration, and spectrophotometry. Our findings support the presence of a preserved and functional visual system in the adult Greenland shark, which seems well-adapted to extreme low-light conditions. Additionally, transcriptomic data suggest a role for DNA repair mechanisms in maintaining retinal integrity over centuries, potentially contributing to the longevity of vision in the longest-living vertebrate.

## Results and Discussion

### Structural and genomic basis of vision in the Greenland shark

Vertebrates typically possess a duplex retina containing both rods and cones. Many deep-sea fish species feature rod-dominated retinas to enhance scotopic vision ^9, 23^. Using ultramicrotomy, we show that the retina of Greenland shark falls into the latter category. Moreover, we found that the Greenland shark retina is characterised by additional dim-light adaptations, such as densely packed and elongated rods, and a high summation between the photoreceptor and the ganglion cell layers (Fig. 1b), similar to what is known from other deep-dwelling or nocturnal sharks ^23-25^. All retinal layers are intact in the Greenland shark, including the photoreceptor layer, the outer nuclear layer, the inner nuclear layer, and the ganglion cell layer (Fig. 1b, Fig. S44). That is, even though all retinas inspected were from adult Greenland sharks estimated to be centuries old, there were no signs of retinal degeneration. This is remarkable given that retinal degeneration is commonly observed in ageing animals, including other long-lived species such as humans ^26, 27^, and suggests that the retinal neurons of the Greenland shark are among the longest-living in any vertebrate.

Vertebrate photoreception relies on a cascade of biochemical reactions mediated by numerous phototransduction genes, many of which are specialised for either rods, cones, or non-visual photoreceptive cells, such as intrinsically photosensitive retinal ganglion cells (ipRGCs) ^8, 28^. Deep-dwelling fish species typically rely more on scotopic vision and, hence, the rod-specific phototransduction genes ^12, 29, 30^. To examine the phototransduction gene repertoire of the Greenland shark, we generated a draft genome for this species and screened it for phototransduction genes. We retrieved functional copies for the full complement of genes required for rod-based phototransduction (Fig. 1c), including *rh1, sag, gnat1, gucy2f, pde6a, pde6b, pde6d, pde6g, grk1, cnga1, gnb5, rcvrn* and *gngt1*. The exact same set of rod-specific phototransduction genes was found in the genomes of five other shark species (*Scyliorhinus canicula, S. torazame, Isurus oxyrinchus, Rhincodon typus*, and *Chiloscyllium plagiosum*; Fig. 1c; Fig. S1), which typically inhabit the upper pelagic zone of tropical to warm temperate seas ^31-35^. Unlike in the other shark genomes inspected, only one functional cone-specific phototransduction gene (*grk7*) could be retrieved from our Greenland shark draft genome. The remaining cone-specific phototransduction genes present in the other sharks were either not found in our draft genome (*arrc*) or were pseudogenised (*rh2, gnat2, pde6c, cnga3*, and *cngb3*) (Fig. 1c). This suggests that Greenland sharks rely on rod-based vision, just like another deep-dwelling shark, the lanternshark *Etmopterus spinax* ^36^ and many deep-sea teleosts ^9^.

We further found that the Greenland shark has a single functional copy of the visual opsin gene *rh1*, and the non-visual opsins *opn3, opn5, rrh* and *va*, similar to other sharks ^37,38^. Like most other shark species, the Greenland shark lacks a copy of the green-sensitive cone opsin gene (*rh2*), and this gene also shows signs of pseudogenization in the Greenland shark genome. Unique among sharks – but similar to what has happened in some cavefish species ^17^ – is the pseudogenization of the non-visual opsin gene *opn4* in the Greenland shark. This gene, also known as melanopsin, mediates light-dependent entrainment of the circadian rhythm in vertebrates ^39^. It thus appears that Greenland sharks may not rely on light to regulate its circadian rhythm or may use alternative mechanisms to mediate this process. We note, however, that the Greenland shark does not exhibit any obvious circadian rhythm in its daily vertical movement pattern ^40, 41^.

Finally, by computing the ω ratio (*i*.*e*., dN/dS) and conducting selection tests, we found that some of the rod-specific phototransduction genes, including *rh1*, showed evidence of relaxed selection relative to other sharks, suggesting that they are under less pressure to be maintained in the Greenland shark genome compared to other species (Fig. 1e; Fig. S12-S42). However, no loss-of-function mutations were detected in this pathway, which would be expected if they were in the process of being pseudogenised ^17^. Furthermore, some rod genes (*e*.*g*., *gnat1* and *sag*) showed evidence of diversifying positive selection, which may suggest adaptive changes related to the optimisation of vision in the Greenland shark.

### Ultrastructural adaptations and circuitry of the Greenland shark retina

To investigate whether the retinal tissue of the Greenland shark actively maintains nuclear organization, which is indicative of active transcription and hence cellular metabolism, we assessed the presence of histone modifications associated with active (H3K27Ac) and repressive (H4K20me3) chromatin states ^42^. We found clear signals for both histone markers in all nuclear layers of the retina (Fig. 2a), indicating active maintenance of chromatin structure across all cell types. Notably, in rod cells, the H3K27Ac marker was primarily localized to the centre of the nucleus, while the repressive H4K20me3 marker was predominantly found near the nuclear lamina (Fig. 2b, 2c). This nuclear organization pattern has previously been associated with diurnality in mammals ^43^. That we now also found this pattern in a species that is predominantly active in dim light suggests that the connection between nuclear organisation and diel activity period is not generalisable in more complex ecological scenarios. Overall, the presence of both active and repressive chromatin states suggests that the retina of the Greenland shark is indeed transcriptionally active.

**Fig. 2.**
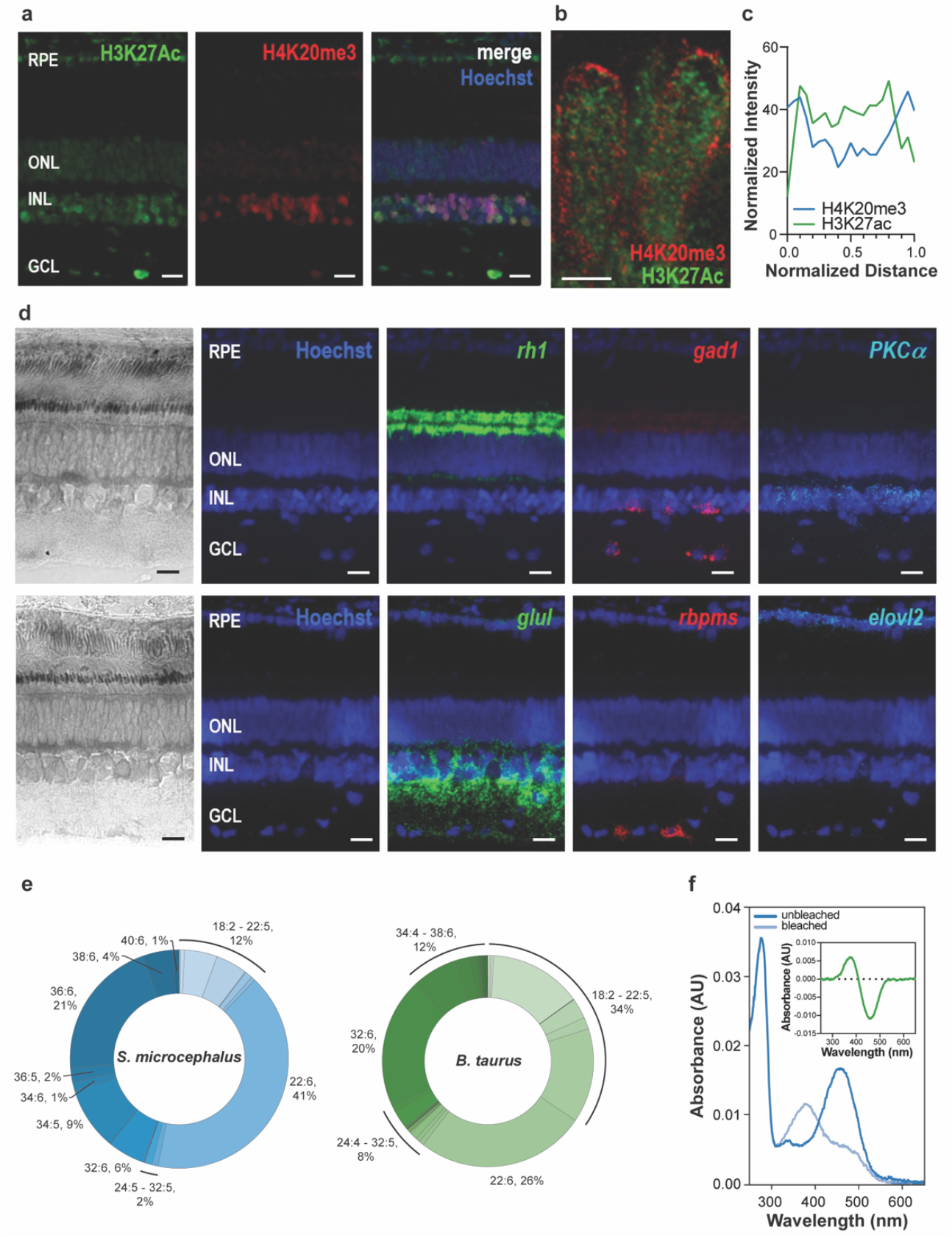
Intact visual circuit and ultrastructural visual adaptations in the Greenland shark. **a**. Immunofluorescent staining of retinal cross-sections from the Greenland shark labelling active (H3K27Ac) and repressed (H4K20me3) chromatin, counter-stained with Hoechst. **b**. Higher magnification image of H4K20me3 and H3K27Ac localization in photoreceptor nuclei. **C**. Distribution of H4K20me3 and H3K27Ac immunofluorescent signal intensity in the nucleus. **d**. RNAscope of key retinal cell markers in retinal cross-sections. Note that the expression of *Elovl2*, a key gene in synthesis of VLC-PUFAs, is confined to the retinal pigment epithelium cells, in contrast to zebrafish expression in Muller glia ^50^ and human expression in cones ^46^. **e**. PUFA composition of bovine and Greenland shark retinas (*n*=2). **f**. Spectral sensitivity of the Greenland shark rhodopsin. Inset depicts the difference (between bleached and unbleached) spectra. ***Scale bars:*** 25 µm **(a, d)**, 5 µm (**b**). ***Abbreviations***: PRL, photoreceptor layer; RPE, retinal pigment epithelium; ONL, outer nuclear layer; INL, inner nuclear layer; GCL, ganglion cell layer.

To assess the integrity of the visual circuit, we employed cell-type specific probes in fluorescent RNA in situ hybridization (RNAscope) to target specific retinal cell types, including rods (using a probe for *rh1*), GABAergic amacrine cells (*gad1*), rod bipolar cells (*pkcα*), Müller glia (*glul*), retinal ganglion cells (*rbpms*) and glycinergic amacrine cells (*slc6a9*) ^44, 45^. Using this approach, we confirmed the presence of all key cell populations necessary for rod-based vision, confirming the integrity of the retinal circuitry in the Greenland shark (Fig. 2d).

Next, we examined the polyunsaturated fatty acid (PUFA) composition of the Greenland shark retina, with a particular focus on omega-3 docosahexaenoic acid (DHA, 22:6n-3) and very long-chain PUFAs (VLC-PUFAs) containing 24 or more carbon atoms. These lipids support rhodopsin function via membrane fluidity and pigment packing ^46-48^ and have been associated with counteracting cold-induced membrane rigidity ^49^. First, we noted that *Elovl2*, a key gene in synthesis of VLC-PUFAs in zebrafish ^50^ and humans ^46^, is also expressed in the Greenland shark retina (Fig. 2d). Targeted mass spectrometry revealed that the retina of the Greenland shark contains an exceptionally high proportion of DHA (41%) compared to the bovine retina (26%) (Fig. 2e), and that VLC-PUFAs constituted 45% of total retinal lipids in the Greenland shark, compared to 35% in mammals. Moreover, the dominant VLC-PUFAs of the Greenland shark featured longer carbon chains (36 carbons) compared to the bovine ones (32 carbons; Fig. 2e). The higher proportion of PUFAs and the longer carbon chains in the Greenland shark compared to other vertebrates might represent yet another adaptation to the deep-sea environment.

Finally, in vertebrates, the spectral sensitivity of opsins is typically tuned to the prevailing wavelengths of light in the environment ^14^. This is also true for the Greenland shark. Spectroscopic analysis of purified, *in vitro* expressed Greenland shark rhodopsin revealed a maximum absorbance wavelength (λ_max_) of 458 nm (Fig. 2), which is shorter compared to most shallow-dwelling sharks ^51^. Short-wavelength shifting of the rhodopsin λ_max_ is a typical adaptation found in deep-sea fishes ^29^, suggesting adaptive evolution of the Greenland shark rhodopsin for life in the deep sea.

### Retinal gene expression in the Greenland Shark

Given that we found a fully intact phototransduction gene repertoire and cell circuitry for rod-based vision in the Greenland shark, we sought to determine whether the expression levels of the phototransduction genes were comparable to those in other shark species and thus, biologically relevant. To address this, we sequenced bulk retinal transcriptomes from the Greenland shark and compared them to publicly available transcriptomic data from retinas of five representative shark species: *S. canicula, S. torazame, Isurus oxyrinchus, R. typus*, and *C. plagiosum*. Our analysis revealed that key rod phototransduction genes were expressed at biologically relevant levels in the retina of the Greenland shark, including rhodopsin (*rh1*), rhodopsin kinase (*grk1*), arrestin (*sag*), transducins (*gnat1* and *gngt1*), and phosphodiesterases (*pde6a, pde6b*, and *pde6g*) (Fig. 3). Expression levels of these genes were found to be comparable to those observed in adult specimens of other shark species (*S. canicula, S. torazame*, and *C. plagiosum*) and higher than those in species sampled as juveniles (*I. oxyrinchus* and *R. typus*).

**Fig. 3.**
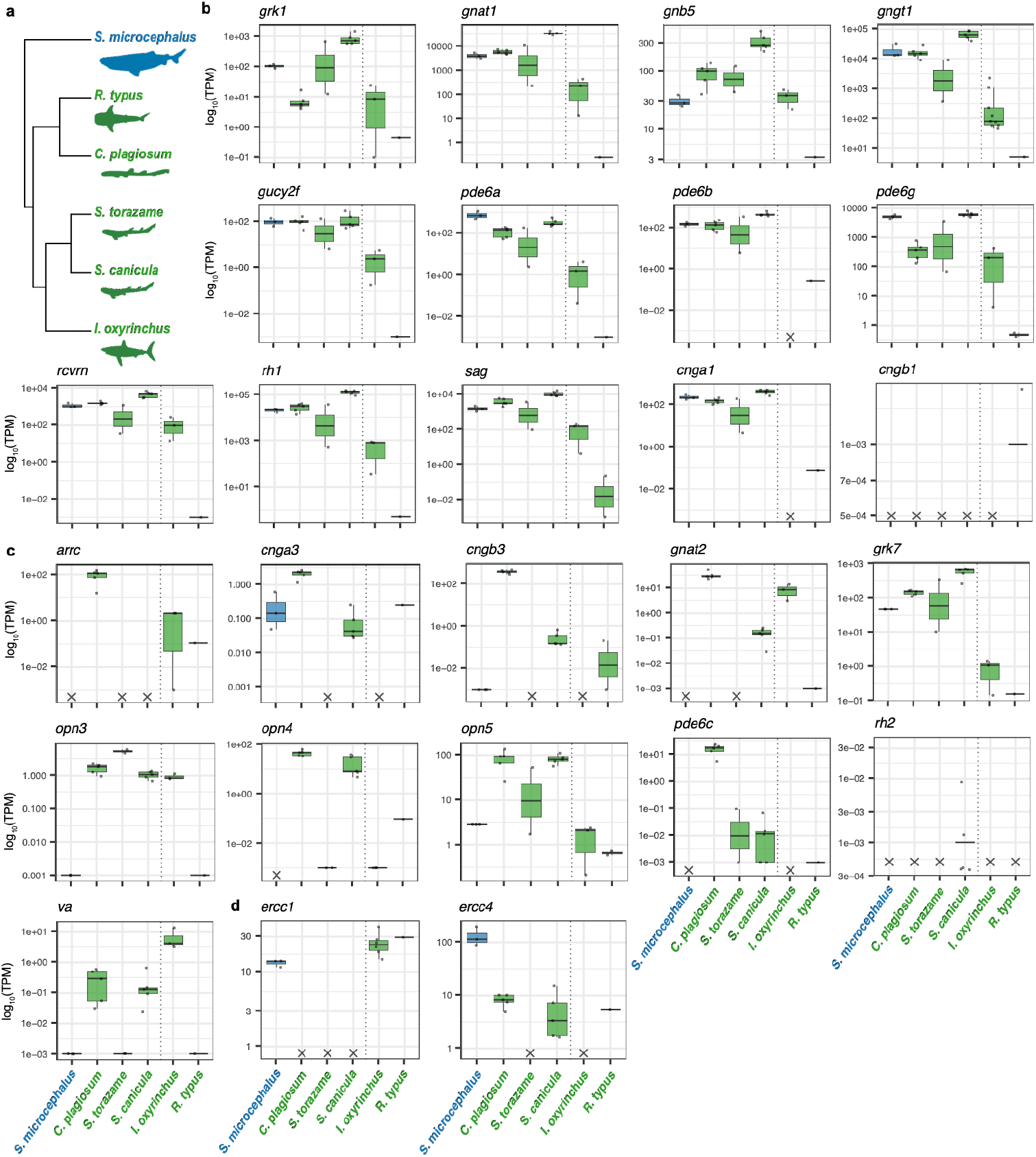
Transcriptomic basis of vision in the Greenland shark. **a**. Phylogeny of species used for gene expression analyses. **b-d**. Retinal expression of genes involved in rod-based phototransduction (**b**), cone-based or non-visual photoreception (**c**) and DNA repair (**d**) in *Somniosus microcephalus* (blue) and five comparison species (green; *Scyliorhinus canicula, S. torazame, Isurus oxyrinchus, Rhincodon typus*, and *Chiloscyllium plagiosum*). Dotted line demarcates data derived from transcriptomes from adult (left) and juvenile (right) specimens. Genes which were not retrieved or pseudogenised are marked with a cross. Note that some genes given in **(b)** are involved in all types of phototransduction (*i*.*e*., *rvcrn, grk1, gnb5* and *gucy2f*).

In contrast, we observed either no expression or low expression of cone phototransduction genes in the Greenland shark (Fig. 3), consistent with the species’ reliance on scotopic vision. Interestingly, the juveniles from our comparison group also exhibited relatively lower levels of most phototransduction genes compared to their adult conspecifics, supporting previous findings suggesting that rod-based phototransduction is not the predominant form of photoreception in the juveniles of some shark species ^52^. Overall, our findings suggest that the Greenland shark preferentially relies on rod-based phototransduction for vision. The transcriptomic basis for scotopic vision in this species is fully intact, comparable to other elasmobranchs, and well-suited to its dim light environment.

Lastly, we investigated whether an efficient DNA repair mechanisms may contribute to the exceptional integrity and sustained retinal function in the retina of the Greenland shark. Specifically, we focused on the ERCC1-XPF DNA repair complex which is known to have an important role in supporting retinal health ^53^. We found that shark species with the longest lifespans, including the Greenland shark, retain the *ercc1* gene, while shorter-lived sharks lacked this gene ^54-59^, and that the Greenland shark exhibits elevated expression of *ercc4 (xpf)* compared to other sharks (Fig. 3). This suggests a robust DNA repair system that may help preserve retinal integrity and function over the extremely long lifespan of the Greenland shark.

### Conclusions

Our findings provide compelling evidence that the Greenland shark (*S. microcephalus*) retains functional vision, despite extreme longevity and an environment characterized by minimal light. The rod phototransduction pathway remains intact, and the loss or pseudogenisation of most cone pathway genes strongly suggests a reliance on scotopic vision. Active transcription of rod phototransduction pathway genes, supported by RNA-seq, RNAscope and chromatin staining, indicates that the retina is functionally preserved.

Furthermore, key visual adaptations revealed by histological analyses and *in vitro* opsin expression are well-aligned with the deep-sea ecology of this species, including elongated, densely packed rods, high rod-to-ganglion cell layer summation and a short-wavelength shift in rhodopsin sensitivity. The absence of retinal degeneration in centuries-old individuals, alongside the preferential retention and elevated expression of DNA repair genes linked to retinal degeneration (*ercc1, ercc4*), suggests a potential mechanism underpinning their long retinal health span. Together, these findings highlight the extraordinary adaptability of vertebrate sensory systems in extreme environments and the remarkable preservation of organ function over hundreds of years.

## Acknowledgements

We would like to thank the crew of the RV Porsild as well as the staff at the Arctic Station in Qeqertarsuaq, Greenland for invaluable help fishing for Greenland sharks. We thank David Salom for his technical expertise with the rhodopsin spectral sensitivity experiment. We acknowledge the staff of the Department of Biosystems Science and Engineering, ETH Zurich for genome and transcriptome sequencing. Panels in some figures were created using BioRender. All genomic and transcriptomic computations were performed at sciCORE (http://scicore.unibas.ch/), the center of scientific computing at University of Basel (with support by the SIB/Swiss Institute of Bioinformatics).

## Data Availability

Newly identified coding sequences (PV442159-PV442194) as well as raw genomic and transcriptomic reads and genome assembly are available through GenBank (PRJNA1246101) and the SRA archive (see Table S2 for accession numbers). All other data are available via Dryad or are provided in the main manuscript or Supplementary Information.

## Methods

### Animal collection and tissue preservation

The Greenland sharks used in this study were caught between 2020 and 2024 using scientific long lines off the coast of the University of Copenhagen’s Arctic Station on Disko Island, Greenland (69°15’N, 53°34’W). All sampling was carried out in accordance with laws and regulations under a permit to collect Greenland sharks for scientific purposes. The work was carried out with authorisation from the Government of Greenland under permits (2020-26794, 2022-24744, 2023-6108, 2024-119) from the Ministry of Fisheries, Hunting & Agriculture and a non-exclusive licence (G24-051) for the utilization of Greenland genetic resources issued by the Ministry of Foreign Affairs, Business and Trade. Individuals were euthanized immediately after capture by direct spinal cord transection. Total body length was measured, and eyecups were dissected and either fixed whole in 4% paraformaldehyde [PFA; 4% (w/v) PFA in 0.01M phosphate-buffered saline] or the retina was removed and fixed in RNAlater or 100% ethanol. Details of all individuals can be found in Table S1.

### Retinal histology

Retinal morphology was assessed for one PFA-fixed eye each from three adult specimens of *S. microcephalus*. Whole, enucleated eyes were post-fixed in 2.5% glutaraldehyde and 2% PFA in 0.1M PBS, progressively dehydrated in increasing concentrations of ethanol, infiltrated with EMbed-812 resin and polymerized at 60°C for 48 h. For light micrographs, 1 μm-thick radial sections were cut on a Reichert-Jung Ultracut E ultramicrotome, deplastified, stained with epoxy tissue stain (Electron Microscopy Sciences; toluidine blue and basic fuchsin), and imaged under brightfield illumination on a Leica DM4 microscope.

### Whole-genome sequencing

A draft genome was sequenced for the Greenland shark, *S. microcephalus*. Briefly, DNA was extracted from frozen tissue from two adult retinas using Zymo Quick-DNA Miniprep Plus kit. DNA quality was assessed on an Agilent TapeStation and library preparation and sequencing was performed by the Department of Biosystems Science and Engineering (ETH Zurich). Individual libraries were prepared using NEBNext Ultra II FS DNA Library Prep kit (Illumina) with 150-bp insert size, pooled, and sequenced as 150-bp paired-end reads on an Illumina NovaSeq6000 with an SP flow cell (300 cycles). Quality control and adapter removal were performed with fastp (v.0.23.4) ^60^, filtered reads were assembled using ABySS (v.2.3.1) using a k-mer size of 80 and k-mer coverage threshold of two ^61^ and the completeness of the assembly was assessed using BUSCO (v.5.4.5) using the vertebrata odb10 database ^62^ (see Table S3 for assembly statistics).

### Chondrichthyes phylogeny

A species tree was constructed for 17 species in the class, Chondrichthyes (see Table S2 for accession numbers). All genome assemblies available for Chondrichthyes were downloaded from the NCBI Genome database. Completeness of each genome was assessed using BUSCO (v.5.4.5) using the vertebrata odb10 database and assemblies with completeness scores over 80% were retained (14 species). An additional three species were included in the phylogeny: *S. microcephalus* and another two species which were used for phototransduction gene mining and expression analyses (*S. torazame*, 76.0% complete; *R. typus*, 75.6% complete). All single-copy orthologs (obtained from BUSCO analysis) present in at least half of the species were aligned using MUSCLE (v.5.1) ^63^, alignments trimmed using trimAl (v.1.4.1) ^64^, and maximum likelihood trees were made for each ortholog using IQ-TREE (v.2.0) ^65^. An unrooted species tree was estimated by inputting concatenated gene trees into ASTRAL (v.5.7.8) ^66^, and the tree was dated and rooted using the least-squared method in IQ-TREE and visualised in iTOL (v.6.9.1) ^67^.

### Phototransduction and DNA repair gene mining

Phototransduction and DNA repair genes were mined from the newly generated draft genome assembly for *S. microcephalus* and from published genome assemblies for five other shark species (*S. canicula*, GCF_902713615.1_sScyCan1.1 ^68^; *S. torazame*, GCA_003427355.1_Storazame_v1.0 ^69^, *R. typus*, GCF_021869965.1_sRhiTyp1.1 ^70^; *I. oxyrinchus*, GCA_026770705.1 ^71^; and *C. plagiosum*, GCF_004010195.1 ^72^), which were selected because they also had publicly available RNAseq data. Genome annotations were obtained for all NCBI genomes, except for *I. oxyrinchus*. The *I. oxyrinchus* genome assembly was annotated by inputting publicly available RNAseq data for this species (see Table S2 for NCBI accession numbers), protein annotations for the four other shark species, and Vertebrata orthologs from OrthoDB (v.12.0) ^73^ into BRAKER (v.3) ^74^.

For *S. microcephalus*, phototransduction and DNA repair gene coding sequences (CDS) were mined using a combination of exon prediction and mapping guided by publicly available reference sequences from NCBI. For the exon prediction approach, scaffolds of interest in the genome assembly were identified using TBLASTN (v.2.11.0) ^75^ and EXONERATE (v.2.4.0) ^76^ was used to predict exons on those scaffolds similar to reference sequences. If the exon prediction approach did not yield a CDS, or if the CDS was fragmented, the CDS was retrieved or completed using a mapping approach. Briefly, raw genomic reads were mapped to reference sequences using HISAT2 (v.2.2.1) ^77^ and the consensus sequence was extracted in Geneious Prime (v.2022.2.2; Biomatters Ltd).

Phototransduction and DNA repair genes were also mined from another five shark species with publicly available genome assemblies. For *S. canicula, R. typus* and *C. plagiosum*, functional genome annotations were available and thus, genes were mined by filtering the annotation (GFF) file to keep only matched gene names and AGAT (v.1.4.1) ^78^ was used to extract the CDS for the longest isoform per gene. For the two species without functional genome annotations (*S. torazame* and *I. oxyrinchus*), all genes were mined using the exon prediction approach described above and annotation files were manually edited to incorporate mined genes.

The identity of every gene mined was confirmed phylogenetically. To that end, gene trees for each clade of interest were generated by downloading reference sequences for outgroup species (*Homo sapiens, Mus musculus* and *Danio rerio*) and other chondrichthyan species from NCBI, aligning these with mined genes using MUSCLE, trimming alignments using trimAl, generating maximum likelihood phylogenies using IQ-TREE and rooting in iTOL (Fig. S1-S11).

### Selection tests on phototransduction and DNA repair genes

The selective forces acting on the newly mined genes were assessed for *S. microcephalus*. For each gene clade, the protein sequences of the Greenland shark were aligned with those from other species in Chondrichthyes using MUSCLE (after removing stop codons and frameshifts present in pseudogene sequences). This protein alignment was then converted to a codon alignment using trimAl and used to construct a gene tree for each clade using IQ-TREE. The codon alignment and the corresponding gene tree were then used to compute maximum likelihood estimates of ω with PAML. Three branch models were used: (i) a free-ratio model, which allows a different ω value per branch; (ii) a two-ratio model assuming one ω for the Greenland shark branches and one ω for all other branches; and (iii) a null one-ratio model assuming that every branch has the same ω ratio. For all genes, the best model was always the free-ratio model (Model 1). The choice between Model 2 and Model 3 was always made using the mean of a likelihood ratio test, using the χ2 distribution with one degree of freedom.

Finally, for each gene clade, RELAX ^79^ and aBSREL ^80^ implemented via the HyPhy framework (v.2.5.63) ^81^ to look for signs of relaxed selection or positive selection in the Greenland shark genes. Greenland shark branches were assigned as “test” branches, while all other branches were assigned as foreground branches, in sequential runs of RELAX and aBSREL.

### Rhodopsin absorption spectrum measurements

The peak spectral sensitivity of the *S. microcephalus* rhodopsin (RHO) was assessed using *in vitro* protein regeneration and spectrophotometry. In brief, HEK293S cells were transfected to express Greenland shark RHO modified to contain the 1D4 epitope derived from the C-terminus of bovine rhodopsin (TETSQVAPA). After 48 h, cells were pelleted down for 1D4-affinity chromatography. Immunoaffinity 1D4 resin was prepared by conjugating purified, anti-Rho antibody (1D4) to CNBr-activated Sepharose 4B beads (Cytiva). Pelleted cells were homogenized, and RHO pigments were reconstituted with 40 µM 11-*cis*-retinal for 1 h at room temperature to regenerate RHO from apo-opsin. Then, 10% DDM was added to solubilize the membrane-enriched pellet, followed by 1 h incubation at 4°C and centrifugation (21,300 x g) for 5 min at 4°C. The supernatant was then filtered through a 0.22 µm polyethersulfone membrane and incubated with 250 µL of 1D4-resin for 1 h at 4°C. Rho-1D4-resin mixture was loaded onto a centrifuge column, washed in a buffer containing 50 mM HEPES, pH 7.5, 0.25 M NaCl, and 10% DDM, and eluted with C-terminal nonapeptide (synthesized by GenScript, NJ, USA) overnight at 4°C. Absorption spectra were recorded using a Varian Cary 50 Scan UV-Vis spectrophotometer (Varian Australian Pty Ltd). The unbleached sample was used as blank, after which the sample was bleached for 10 min with a white-light, 875-Lumens bulb. The difference in absorption spectrum was then recorded.

### Nuclear chromatin staining and quantification

Eyecups were fixed in 4% paraformaldehyde overnight, cryoprotected by immersion in a sucrose gradient (10% and 20% sucrose for 1 hour at room temperature, and 30% sucrose overnight at 4 °C), embedded in Tissue-Tek OCT (Sakura, Torrance, CA) and frozen on a conductive metal block placed on dry ice. After cryosectioning, sections were blocked in 5% BSA, 0.3% TritonX-100 for 1.5 hour at room temperature to minimize nonspecific binding. Sections were then incubated with primary antibodies (mouse anti-trimethyl histone H4 (sc-134216), 1:200 and rabbit anti-H3K27ac (ab-479), 1:200) diluted in 5% BSA, 0.1% TritonX-100 overnight at 4°C. Following three washes with PBS, sections were incubated with fluorescently-labeled secondary antibodies diluted in 5% BSA, 0.1% TritonX-100 for 1 hour at room temperature. Following three washes with PBS, nuclei were counterstained with Hoescht 33342 (Thermo), and sections were mounted using ProLong Gold Antifade (Thermo). Immunostained sections were imaged on a Zeiss LSM900 confocal microscope with Airyscan 2 at 40X magnification. Quantification of nuclear chromatin distribution was performed using ImageJ. For each cell, two straight lines were drawn across the nucleus, and intensity profiles for the H4K20me3, H3K27ac, and Hoechst channels were measured. The intensity profiles were normalized within each channel and plotted against the normalized distance along each line.

### RNAscope

In situ hybridization was performed using the RNAscope^®^ Multiplex Fluorescent Assay v2 (ACD Diagnostics) following modifications as described previously ^82^. Briefly, frozen histologic sections of fixed shark eyes were pretreated per manual using hydrogen peroxide and target retrieval reagents including protease IV. Probes were then hybridized according to the protocol and then detected with TSA Plus^®^ Fluorophores fluorescein, cyanine 3, and cyanine 5. Sections were mounted with Prolong Gold Antifade (Thermo Fisher) with coverslip for imaging and imaged (Keyence BZ-X700). Probes specific for mouse and human transcripts were designed by the manufacturer (see Table S4).

### Lipidomic analysis

Lipid purification and fatty acid were performed as previously described ^83^. Briefly, the tissue was homogenized and lipids were extracted as in ^84^, and dried under nitrogen, and stored at - 20 °C until subsequent lipid analysis. The total fatty acids were released through acid hydrolysis and extracted by hexane then, the sample was evaporated under nitrogen and stored at -20 °C until subsequent lipid analysis. Lipidomic and fatty acid analyses were performed on Q Exactive mass spectrometer coupled with a Vanquish liquid chromatography system (Thermo Fisher Scientific) using an ESI spray source. Quantification was performed relative to the total intensity of lipids or PUFAs. Data visualization was performed on Prism 7 software (GraphPad Software, Inc.).

### Retinal transcriptome sequencing

Retinal transcriptomes were sequenced for three individuals for *S. microcephalus*. Briefly, total RNA was extracted from one RNAlater-fixed retina from each of three adult sharks using the QuickRNA Miniprep kit (Zymo). RNA quality was assessed on an Agilent TapeStation and library preparation and sequencing was performed by the Department of Biosystems Science and Engineering (ETH Zurich). Individual libraries were prepared using TruSeq stranded total RNA ribo-zero gold kit (Illumina), pooled, and sequenced as 100-bp paired-end reads using BRAVO Sequencing on an Illumina NovaSeq6000 with an S4 flow cell (200 cycles). Quality control and adapter removal was performed with fastp (v.0.23.4).

### Phototransduction and DNA repair gene expression

Phototransduction and DNA repair gene expression was quantified for the Greenland shark and five other shark species for which retinal transcriptome data were available on NCBI (see Table S2 for accession numbers). For the five other species, transcriptomes were pre-processed using fastp and mapped to annotated genomes using STAR (v.2.7.10b) ^85^ with -- outFilterMultimapNmax 1 --outFilterMatchNminOverLread 0.4 -- outFilterScoreMinOverLread 0.4, and filtered out mapped singletons for paired-end data.

Read counts were performed using the HTSeq-count script from the HTSeq framework (v.2.0.2) ^86^ and were used to calculate TPM values for all genes. For the genes of interest, TPM values were plotted against the proportion of reads mapped (calculated as the number of reads mapped divided by the total number of reads in the transcriptome) in R (v.4.4.0) ^87^ and a linear regression was performed to generate an equation that describes the relationship between the two variables for each gene (Fig. S43).

For the Greenland shark, an annotation file was manually generated for all genes that were mined and this was used to guide mapping against those sequences using STAR. Similar to the other species, mapped singletons were filtered out and read counts were performed using HTSeq-count. The equations generated using the other five shark species were used to extrapolate TPM values from the proportion of mapped reads for each gene expressed in the Greenland shark.

## References

1. Nielsen, J., R. Hedeholm, P. Bushnell, R. Brill, J. Olsen, J. Heinemeier, J. Christiansen, M. Simon, K. Steffensen and J. Steffensen. (2016) Eye lens radiocarbon reveals centuries of longevity in the Greenland shark (Somniosus microcephalus). Science. 353: 702.

2. Nielsen, J., R.B. Hedeholm, M. Simon and J.F. Steffensen. (2014) Distribution and feeding ecology of the Greenland shark (Somniosus microcephalus) in Greenland waters. Polar Biology. 37(1): 37–46.

3. Mecklenburg, C., A. Lynghammar, E. Johannesen, I. Byrkjedal, J. Christiansen, A. Dolgov, O. Karamushko, T. Mecklenburg, P. Moller, D. Steinke and R. Wienerroither. (2018) Marine Fishes of the Arctic Region. Conservation of Arctic Flora and Fauna. Vol. 1. Akureyri, Iceland.

4. MacNeil, M.A., B.C. McMeans, N.E. Hussey, P. Vecsei, J. Svavarsson, K.M. Kovacs, C. Lydersen, M.A. Treble, G.B. Skomal, M. Ramsey and A.T. Fisk. (2012) Biology of the Greenland shark Somniosus microcephalus. Journal of Fish Biology. 80(5): 991–1018.

5. Berland, B. (1961) Copepod Ommatokoita elongata (Grant) in the Eyes of the Greenland Shark—a Possible Cause of Mutual Dependence. Nature. 191(4790): 829–830.

6. Benz, G.W., J.D. Borucinska, L.F. Lowry and H.E. Whiteley. (2002) Ocular lesions associated with attachment of the copepod Ommatokoita elongata (Lernaeopodidae: Siphonostomatoida) to corneas of Pacific sleeper sharks Somniosus pacificus captured off Alaska in Prince William Sound. J Parasitol. 88(3): 474–81.

7. Lamb, T.D. (2016) Why rods and cones? Eye (Lond). 30(2): 179–85.

8. Lamb, T.D. (2013) Evolution of phototransduction, vertebrate photoreceptors and retina. Prog Retin Eye Res. 36: 52–119.

9. de Busserolles, F., L. Fogg, F. Cortesi and J. Marshall. (2020) The exceptional diversity of visual adaptations in deep-sea teleost fishes. Seminars in Cell & Developmental Biology. 106: 20–30.

10. Fogg, L.G., F. Cortesi, D. Lecchini, C. Gache, N.J. Marshall and F. de Busserolles. (2022) Development of dim-light vision in the nocturnal reef fish family Holocentridae I: Retinal gene expression. Journal of Experimental Biology. 225(17): jeb244513.

11. Fogg, L.G., F. Cortesi, D. Lecchini, C. Gache, N.J. Marshall and F. de Busserolles. (2022) Development of dim-light vision in the nocturnal reef fish family Holocentridae II: Retinal morphology Journal of Experimental Biology. 225(17): jeb244740.

12. Fogg, L.G., S. Isari, J.E. Barnes, J.S. Patel, N.J. Marshall, W. Salzburger, F. Cortesi and F. de Busserolles. (2024) Deep-sea fish reveal alternative pathway for vertebrate visual development. bioRxiv: 2024.10.10.617579.

13. Cortesi, F., L.J. Mitchell, V. Tettamanti, L.G. Fogg, F. de Busserolles, K.L. Cheney and N.J. Marshall. (2020) Visual system diversity in coral reef fishes. Seminars in Cell & Developmental Biology. 106: 31–42.

14. Musilova, Z., W. Salzburger and F. Cortesi. (2021) The Visual Opsin Gene Repertoires of Teleost Fishes: Evolution, Ecology, and Function. Annual Review of Cell and Developmental Biology. 37(1): 441–468.

15. Hauzman, E. (2020) Adaptations and evolutionary trajectories of the snake rod and cone photoreceptors. Seminars in Cell & Developmental Biology. 106: 86–93.

16. de Busserolles, F. and N.J. Marshall. (2017) Seeing in the deep-sea: visual adaptations in lanternfishes. Philos Trans R Soc Lond B Biol Sci. 372(1717).

17. Policarpo, M., J. Fumey, P. Lafargeas, D. Naquin, C. Thermes, M. Naville, C. Dechaud, J.-N. Volff, C. Cabau, C. Klopp, P.R. Møller, L. Bernatchez, E. García-Machado, S. Rétaux and D. Casane. (2020) Contrasting gene decay in subterranean vertebrates: insights from cavefishes and fossorial mammals. Molecular Biology and Evolution.

18. Simon, N., S. Fujita, M. Porter and M. Yoshizawa. (2019) Expression of extraocular opsin genes and light-dependent basal activity of blind cavefish. PeerJ. 7: e8148.

19. Protas, M. and W.R. Jeffery. (2012) Evolution and development in cave animals: from fish to crustaceans. WIREs Developmental Biology. 1(6): 823–845.

20. Edwards, J.E., E. Hiltz, F. Broell, P.G. Bushnell, S.E. Campana, J.S. Christiansen, B.M. Devine, J.J. Gallant, K.J. Hedges, M.A. MacNeil, B.C. McMeans, J. Nielsen, K. Præbel, G.B. Skomal, J.F. Steffensen, R.P. Walter, Y.Y. Watanabe, D.L. VanderZwaag and N.E. Hussey. (2019) Advancing Research for the Management of Long-Lived Species: A Case Study on the Greenland Shark. Frontiers in Marine Science. 6.

21. Yopak, K.E., B.C. McMeans, C.G. Mull, K.W. Feindel, K.M. Kovacs, C. Lydersen, A.T. Fisk and S.P. Collin. (2019) Comparative Brain Morphology of the Greenland and Pacific Sleeper Sharks and its Functional Implications. Sci Rep. 9(1): 10022.

22. Bartas, M., J. Cerven, N. Valková, A. Volná, M. Dobrovolna, L. Šislerová, H. Baldvinsson, P. Pecinka and V. Brázda. (2024) RNA analysis of the longest living vertebrate Greenland shark revealed an abundance of LINE-like elements in its transcriptome. Czech Polar Reports. 13.

23. Newman, A.S., J.N. Marshall and S.P. Collin. (2013) Visual Eyes: A Quantitative Analysis of the Photoreceptor Layer in Deep-Sea Sharks. Brain Behavior and Evolution. 82(4): 237–249.

24. Peel, L.R., S.P. Collin and N.S. Hart. (2020) Retinal topography and spectral sensitivity of the Port Jackson shark (Heterodontus portusjacksoni). Journal of Comparative Neurology. 528(17): 2831–2847.

25. Schieber, N.L., S.P. Collin and N.S. Hart. (2012) Comparative retinal anatomy in four species of elasmobranch. Journal of Morphology. 273(4): 423–440.

26. Jackson, G.R., C. Owsley and C.A. Curcio. (2002) Photoreceptor degeneration and dysfunction in aging and age-related maculopathy. Ageing research reviews. 1(3): 381–396.

27. Guymer, R.H. and T.G. Campbell. (2023) Age-related macular degeneration. The Lancet. 401(10386): 1459–1472.

28. Do, M.T.H. (2019) Melanopsin and the Intrinsically Photosensitive Retinal Ganglion Cells: Biophysics to Behavior. Neuron. 104(2): 205–226.

29. Musilova, Z., F. Cortesi, M. Matschiner, W.I.L. Davies, J.S. Patel, S.M. Stieb, F. de Busserolles, M. Malmstrom, O.K. Torresen, C.J. Brown, J.K. Mountford, R. Hanel, D.L. Stenkamp, K.S. Jakobsen, K.L. Carleton, S. Jentoft, J. Marshall and W. Salzburger. (2019) Vision using multiple distinct rod opsins in deep-sea fishes. Science. 364(6440): 588–592.

30. Lupše, N., F. Cortesi, M. Freese, L. Marohn, J.-D. Pohlmann, K. Wysujack, R. Hanel and Z. Musilova. (2021) Visual Gene Expression Reveals a cone-to-rod Developmental Progression in Deep-Sea Fishes. Molecular biology and evolution. 38(12): 5664–5677.

31. Bacchet, P., T. Zysman and Y. Lefevre. (2016) Guide des poissons de Tahiti et ses iles. 4th ed. Tahiti: Au vent des iles.

32. Weigmann, S. (2016) Annotated checklist of the living sharks, batoids and chimaeras (Chondrichthyes) of the world, with a focus on biogeographical diversity. Journal of Fish Biology. 88(3): 837–1037.

33. Bianchi, G., K.E. Carpenter, J.-P. Roux, F.J. Molloy, D. Boyer and H.J. Boyer. (1999) Field guide to the living marine resources of Namibia. FAO species identification guide for fishery purposes. . Rome: FAO.

34. Capapé, C., Y. Vergne, C. Reynaud, O. Guélorget and J. Quignard. (2008) Maturity, fecundity and occurrence of the smallspotted catshark Scyliorhinus canicula (Chondrichthyes: Scyliorhinidae) off the Languedocian coast (southern France, north-western Mediterranean). Vie et Milieu/Life & Environment: 47–55.

35. Ito, N., M. Fujii, K. Nohara and S. Tanaka. (2022) Scyliorhinus hachijoensis, a new species of catshark from the Izu Islands, Japan (Carcharhiniformes: Scyliorhinidae). Zootaxa. 5092(3): 331–349.

36. Delroisse, J., L. Duchatelet, P. Flammang and J. Mallefet. (2018) De novo transcriptome analyses provide insights into opsin-based photoreception in the lanternshark Etmopterus spinax. PLoS One. 13(12): e0209767.

37. Yamaguchi, K., M. Koyanagi and S. Kuraku. (2021) Visual and nonvisual opsin genes of sharks and other nonosteichthyan vertebrates: Genomic exploration of underwater photoreception. J Evol Biol. 34(6): 968–976.

38. Hart, N.S. (2020) Vision in sharks and rays: Opsin diversity and colour vision. Seminars in Cell & Developmental Biology. 106: 12–19.

39. Pan, D., Z. Wang, Y. Chen and J. Cao. (2023) Melanopsin-mediated optical entrainment regulates circadian rhythms in vertebrates. Communications Biology. 6(1): 1054.

40. Fisk, A.T., C. Lydersen and K.M. Kovacs. (2012) Archival pop-off tag tracking of Greenland sharks Somniosus microcephalus in the High Arctic waters of Svalbard, Norway. Marine Ecology Progress Series. 468: 255–265.

41. Stokesbury, M., C. Harvey-Clark, J. Hay Gallant, B. Block and R. Myers. (2005) Movement and environmental preferences of Greenland sharks (Somniosus microcephalus) electronically tagged in the St. Lawrence Estuary, Canada. Marine Biology. 148: 159–165.

42. Telese, F., A. Gamliel, D. Skowronska-Krawczyk, I. Garcia-Bassets and M.G. Rosenfeld. (2013) “Seq-ing” insights into the epigenetics of neuronal gene regulation. Neuron. 77(4): 606–23.

43. Solovei, I., M. Kreysing, C. Lanctôt, S. Kösem, L. Peichl, T. Cremer, J. Guck and B. Joffe. (2009) Nuclear Architecture of Rod Photoreceptor Cells Adapts to Vision in Mammalian Evolution. Cell. 137(2): 356–368.

44. Chao, D.L. and D. Skowronska-Krawczyk. (2020) ELOVL2: Not just a biomarker of aging. Translational Medicine of Aging. 4: 78–80.

45. Vidal-Vázquez, N., I. Hernández-Núñez, P. Carballo-Pacoret, S. Salisbury, P.R. Villamayor, F. Hervas-Sotomayor, X. Yuan, F. Lamanna, C. Schneider, J. Schmidt, S. Mazan, H. Kaessmann, F. Adrio, D. Robledo, A. Barreiro-Iglesias and E. Candal. (2025) A single-nucleus RNA sequencing atlas of the postnatal retina of the shark Scyliorhinus canicula. Scientific Data. 12(1): 228.

46. Lewandowski, D., C.L. Sander, A. Tworak, F. Gao, Q. Xu and D. Skowronska-Krawczyk. (2022) Dynamic lipid turnover in photoreceptors and retinal pigment epithelium throughout life. Prog Retin Eye Res. 89: 101037.

47. Agbaga, M.-P., D.K. Merriman, R.S. Brush, T.A. Lydic, S.M. Conley, M.I. Naash, S. Jackson, A.S. Woods, G.E. Reid, J.V. Busik and R.E. Anderson. (2018) Differential composition of DHA and very-long-chain PUFAs in rod and cone photoreceptors. Journal of Lipid Research. 59(9): 1586–1596.

48. Sander, C.L., A.E. Sears, A.F.M. Pinto, E.H. Choi, S. Kahremany, F. Gao, D. Salom, H. Jin, E. Pardon, S. Suh, Z. Dong, J. Steyaert, A. Saghatelian, D. Skowronska-Krawczyk, P.D. Kiser and K. Palczewski. (2021) Nano-scale resolution of native retinal rod disk membranes reveals differences in lipid composition. J Cell Biol. 220(8).

49. Winnikoff, J.R., S.H.D. Haddock and I. Budin. (2021) Depth-and temperature-specific fatty acid adaptations in ctenophores from extreme habitats. J Exp Biol. 224(21).

50. Dasyani, M., F. Gao, Q. Xu, D. Van Fossan, E. Zhang, A. F. M. Pinto, A. Saghatelian, D. Skowronska-Krawczyk and D.L. Chao. (2020) Elovl2 Is Required for Robust Visual Function in Zebrafish. Cells. 9(12): 2583.

51. Hart, N.S., T.D. Lamb, H.R. Patel, A. Chuah, R.C. Natoli, N.J. Hudson, S.C. Cutmore, W.I.L. Davies, S.P. Collin and D.M. Hunt. (2019) Visual Opsin Diversity in Sharks and Rays. Molecular Biology and Evolution. 37(3): 811–827.

52. Domingues, R.R., V.A. Mastrochirico-Filho, N.J. Mendes, D.T. Hashimoto, R. Coelho, V.P. da Cruz, A. Antunes, F. Foresti and F.F. Mendonça. (2020) Comparative eye and liver differentially expressed genes reveal monochromatic vision and cancer resistance in the shortfin mako shark (Isurus oxyrinchus). Genomics. 112(6): 4817–4826.

53. Narasimhan, A., S.H. Min, L.L. Johnson, H. Roehrich, W. Cho, T.K. Her, C. Windschitl, R.D. O’Kelly, L. Angelini, M.J. Yousefzadeh, L.K. McLoon, W.W. Hauswirth, P.D. Robbins, D. Skowronska-Krawczyk and L.J. Niedernhofer. (2024) The Ercc1(-/Δ) mouse model of XFE progeroid syndrome undergoes accelerated retinal degeneration. Aging Cell: e14419.

54. Bishop, S., M. Francis, C. Duffy and J. Montgomery. (2006) Age, growth, maturity, longevity and natural mortality of the shortfin mako shark (Isurus oxyrinchus) in New Zealand waters. Marine and Freshwater Research. 57: 143–154.

55. Perry, C.T., J. Figueiredo, J.J. Vaudo, J. Hancock, R. Rees and M. Shivji. (2018) Comparing length-measurement methods and estimating growth parameters of free-swimming whale sharks (<i>Rhincodon typus</i>) near the South Ari Atoll, Maldives. Marine and Freshwater Research. 69(10): 1487–1495.

56. Moreira, I., I. Figueiredo, I. Farias, N. Lagarto, C. Maia, J. Robalo and T. Moura. (2022) Growth and maturity of the lesser-spotted dogfish (Linnaeus, 1758) in the southern Portuguese continental coast. Journal of Fish Biology. 100(1): 315–319.

57. Michael, S.W. (1993) Reef sharks and rays of the world. A guide to their identification, behaviour, and ecology. 2009/05/11 ed. Sea Challengers. Vol. 73. Monterey, California: Journal of the Marine Biological Association of the United Kingdom.

58. Chen, W., P. Chen, K.-M. Liu and S.-B. Wang. (2007) Age and growth estimates of the Whitespotted Bamboo Shark, Chiloscyllium plagiosum, in the Northern Waters of Taiwan. Zoological Studies. 46: 92–102.

59. Fahmi, W. Kurniawan, I.R. Tibbetts, S. Oktaviyani, C.L. Dudgeon and M.B. Bennett. (2021) Age and growth of the tropical oviparous shark, Chiloscyllium punctatum in Indonesian waters. J Fish Biol. 99(3): 921–930.

60. Chen, S., Y. Zhou, Y. Chen and J. Gu. (2018) fastp: an ultra-fast all-in-one FASTQ preprocessor. Bioinformatics. 34(17): i884–i890.

61. Simpson, J.T., K. Wong, S.D. Jackman, J.E. Schein, S.J. Jones and I. Birol. (2009) ABySS: a parallel assembler for short read sequence data. Genome Res. 19(6): 1117–23.

62. Simão, F.A., R.M. Waterhouse, P. Ioannidis, E.V. Kriventseva and E.M. Zdobnov. (2015) BUSCO: assessing genome assembly and annotation completeness with single-copy orthologs. Bioinformatics. 31(19): 3210–3212.

63. Edgar, R.C. (2004) MUSCLE: a multiple sequence alignment method with reduced time and space complexity. BMC Bioinformatics. 5(1): 113.

64. Capella-Gutiérrez, S., J.M. Silla-Martínez and T. Gabaldón. (2009) trimAl: a tool for automated alignment trimming in large-scale phylogenetic analyses. Bioinformatics. 25(15): 1972–3.

65. Minh, B.Q., H.A. Schmidt, O. Chernomor, D. Schrempf, M.D. Woodhams, A. von Haeseler and R. Lanfear. (2020) IQ-TREE 2: New Models and Efficient Methods for Phylogenetic Inference in the Genomic Era. Molecular Biology and Evolution. 37(5): 1530–1534.

66. Zhang, C., M. Rabiee, E. Sayyari and S. Mirarab. (2018) ASTRAL-III: polynomial time species tree reconstruction from partially resolved gene trees. BMC Bioinformatics. 19(6): 153.

67. Letunic, I. and P. Bork. (2021) Interactive Tree Of Life (iTOL) v5: an online tool for phylogenetic tree display and annotation. Nucleic Acids Research. 49(W1): W293–W296.

68. Mayeur, H., J. Leyhr, J. Mulley, N. Leurs, L. Michel, K. Sharma, R. Lagadec, J.-M. Aury, O.G. Osborne, P. Mulhair, J. Poulain, S. Mangenot, D. Mead, M. Smith, C. Corton, K. Oliver, J. Skelton, E. Betteridge, J. Dolucan, O. Dudchenko, A.D. Omer, D. Weisz, E.L. Aiden, S.A. McCarthy, Y. Sims, J. Torrance, A. Tracey, K. Howe, T. Baril, A. Hayward, C. Martinand-Mari, S. Sanchez, T. Haitina, K. Martin, S.I. Korsching, S. Mazan and M. Debiais-Thibaud. (2024) The Sensory Shark: High-quality Morphological, Genomic and Transcriptomic Data for the Small-spotted Catshark Scyliorhinus Canicula Reveal the Molecular Bases of Sensory Organ Evolution in Jawed Vertebrates. Molecular Biology and Evolution. 41(12).

69. Hara, Y., K. Yamaguchi, K. Onimaru, M. Kadota, M. Koyanagi, S.D. Keeley, K. Tatsumi, K. Tanaka, F. Motone, Y. Kageyama, R. Nozu, N. Adachi, O. Nishimura, R. Nakagawa, C. Tanegashima, I. Kiyatake, R. Matsumoto, K. Murakumo, K. Nishida, A. Terakita, S. Kuratani, K. Sato, S. Hyodo and S. Kuraku. (2018) Shark genomes provide insights into elasmobranch evolution and the origin of vertebrates. Nature Ecology & Evolution. 2(11): 1761–1771.

70. Yamaguchi, K., Y. Uno, M. Kadota, O. Nishimura, R. Nozu, K. Murakumo, R. Matsumoto, K. Sato and S. Kuraku. (2023) Elasmobranch genome sequencing reveals evolutionary trends of vertebrate karyotype organization. Genome Res. 33(9): 1527–1540.

71. Stanhope, M.J., K.M. Ceres, Q. Sun, M. Wang, J.D. Zehr, N.J. Marra, A.P. Wilder, C. Zou, A.M. Bernard, P. Pavinski-Bitar, M.G. Lokey and M.S. Shivji. (2023) Genomes of endangered great hammerhead and shortfin mako sharks reveal historic population declines and high levels of inbreeding in great hammerhead. iScience. 26(1): 105815.

72. Zhang, Y., H. Gao, H. Li, J. Guo, B. Ouyang, M. Wang, Q. Xu, J. Wang, M. Lv, X. Guo, Q. Liu, L. Wei, H. Ren, Y. Xi, Y. Guo, B. Ren, S. Pan, C. Liu, X. Ding, H. Xiang, Y. Yu, Y. Song, L. Meng, S. Liu, J. Wang, Y. Jiang, J. Shi, S. Liu, J.S.M. Sabir, M.J. Sabir, M. Khan, N.H. Hajrah, S. Ming-Yuen Lee, X. Xu, H. Yang, J. Wang, G. Fan, N. Yang and X. Liu. (2020) The White-Spotted Bamboo Shark Genome Reveals Chromosome Rearrangements and Fast-Evolving Immune Genes of Cartilaginous Fish. iScience. 23(11): 101754.

73. Kuznetsov, D., F. Tegenfeldt, M. Manni, M. Seppey, M. Berkeley, E.V. Kriventseva and E.M. Zdobnov. (2023) OrthoDB v11: annotation of orthologs in the widest sampling of organismal diversity. Nucleic Acids Res. 51(D1): D445–d451.

74. Gabriel, L., T. Brůna, K.J. Hoff, M. Ebel, A. Lomsadze, M. Borodovsky and M. Stanke. (2024) BRAKER3: Fully automated genome annotation using RNA-seq and protein evidence with GeneMark-ETP, AUGUSTUS, and TSEBRA. Genome Res. 34(5): 769–777.

75. Sayers, E.W., E.E. Bolton, J.R. Brister, K. Canese, J. Chan, D.C. Comeau, R. Connor, K. Funk, C. Kelly, S. Kim, T. Madej, A. Marchler-Bauer, C. Lanczycki, S. Lathrop, Z. Lu, F. Thibaud-Nissen, T. Murphy, L. Phan, Y. Skripchenko, T. Tse, J. Wang, R. Williams, B.W. Trawick, K.D. Pruitt and S.T. Sherry. (2022) Database resources of the national center for biotechnology information. Nucleic Acids Res. 50(D1): D20–d26.

76. Slater, G.S.C. and E. Birney Automated generation of heuristics for biological sequence comparison. BMC bioinformatics, 2005. 6, 31 DOI: 10.1186/1471-2105-6-31.

77. Kim, D., J.M. Paggi, C. Park, C. Bennett and S.L. Salzberg. (2019) Graph-based genome alignment and genotyping with HISAT2 and HISAT-genotype. Nature Biotechnology. 37(8): 907–915.

78. Dainat, J. Another Gtf/Gff Analysis Toolkit (AGAT): Resolve interoperability issues and accomplish more with your annotations. in Plant and Animal Genome XXIX Conference. 2022.

79. Wertheim, J.O., B. Murrell, M.D. Smith, S.L. Kosakovsky Pond and K. Scheffler. (2014) RELAX: Detecting Relaxed Selection in a Phylogenetic Framework. Molecular Biology and Evolution. 32(3): 820–832.

80. Smith, M.D., J.O. Wertheim, S. Weaver, B. Murrell, K. Scheffler and S.L. Kosakovsky Pond. (2015) Less is more: an adaptive branch-site random effects model for efficient detection of episodic diversifying selection. Mol Biol Evol. 32(5): 1342–53.

81. Pond, S.L.K., S.D.W. Frost and S.V. Muse. (2004) HyPhy: hypothesis testing using phylogenies. Bioinformatics. 21(5): 676–679.

82. Xu, Q., C. Rydz, V.A. Nguyen Huu, L. Rocha, C. Palomino La Torre, I. Lee, W. Cho, M. Jabari, J. Donello, D.C. Lyon, R.T. Brooke, S. Horvath, R.N. Weinreb, W.K. Ju, A. Foik and D. Skowronska-Krawczyk. (2022) Stress induced aging in mouse eye. Aging Cell. 21(12): e13737.

83. Gao, F., E. Tom, S.A. Lieffrig, S.C. Finnemann and D. Skowronska-Krawczyk. (2023) A novel quantification method for retinal pigment epithelium phagocytosis using a very-long-chain polyunsaturated fatty acids-based strategy. Front Mol Neurosci. 16: 1279457.

84. Bligh, E.G. and W.J. Dyer. (1959) A rapid method of total lipid extraction and purification. Can J Biochem Physiol. 37(8): 911–7.

85. Dobin, A., C.A. Davis, F. Schlesinger, J. Drenkow, C. Zaleski, S. Jha, P. Batut, M. Chaisson and T.R. Gingeras. (2013) STAR: ultrafast universal RNA-seq aligner. Bioinformatics. 29(1): 15–21.

86. Anders, S., P.T. Pyl and W. Huber. (2015) HTSeq-a Python framework to work with high-throughput sequencing data. Bioinformatics. 31(2): 166–9.

87. R Core Team. (2022) R: A language and environment for statistical computing. R Foundation for Statistical Computing, Vienna, Austria.

