## Supplementary Information for "The visual system of the longest-living vertebrate, the Greenland shark"

### Supplementary Figures and Tables

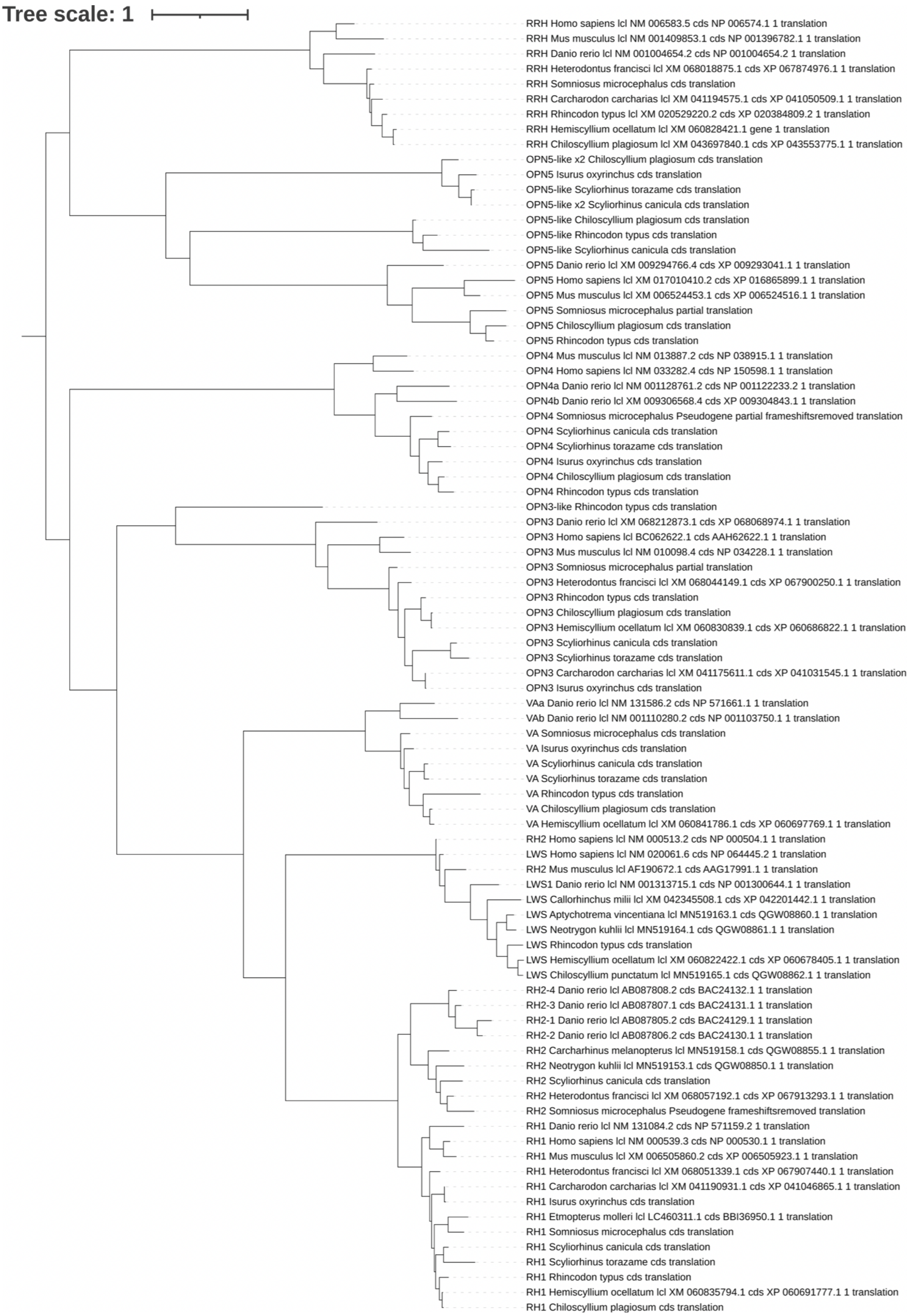


**Fig S1. Opsin phylogeny.** Gene tree with sequences from several shark species and outgroup species (*Danio rerio, Homo sapiens* and *Mus musculus*). The scale bar denotes substitutions per site.

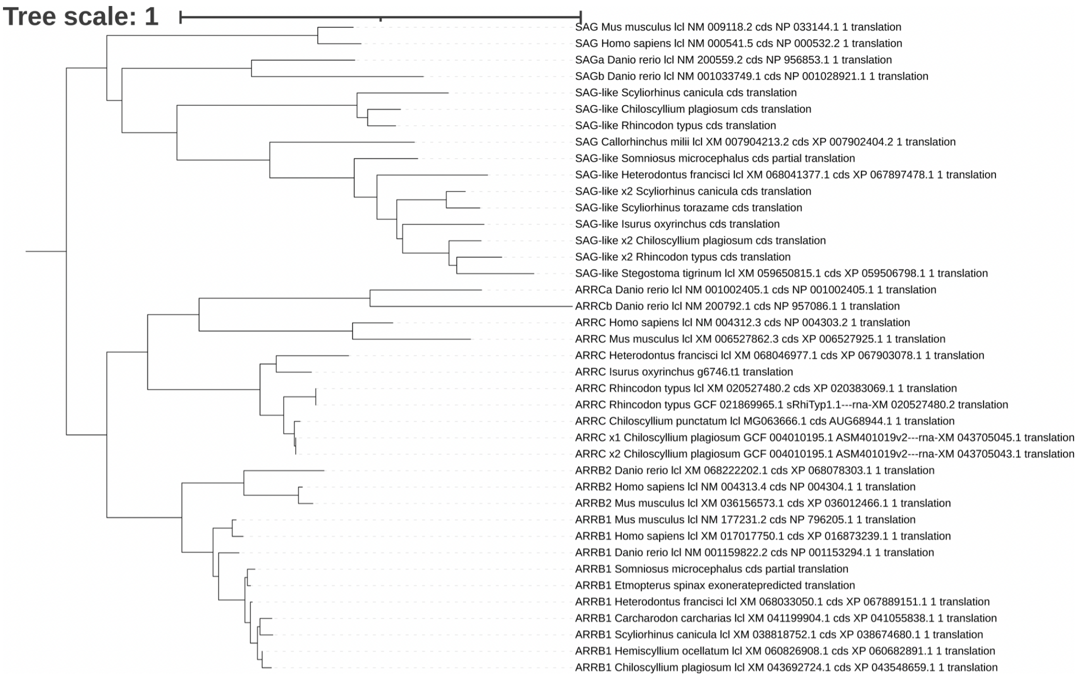


**Fig S2. Arrestin phylogeny.** Gene tree with sequences from several shark species and outgroup species (*Danio rerio, Homo sapiens* and *Mus musculus*). The scale bar denotes substitutions per site.

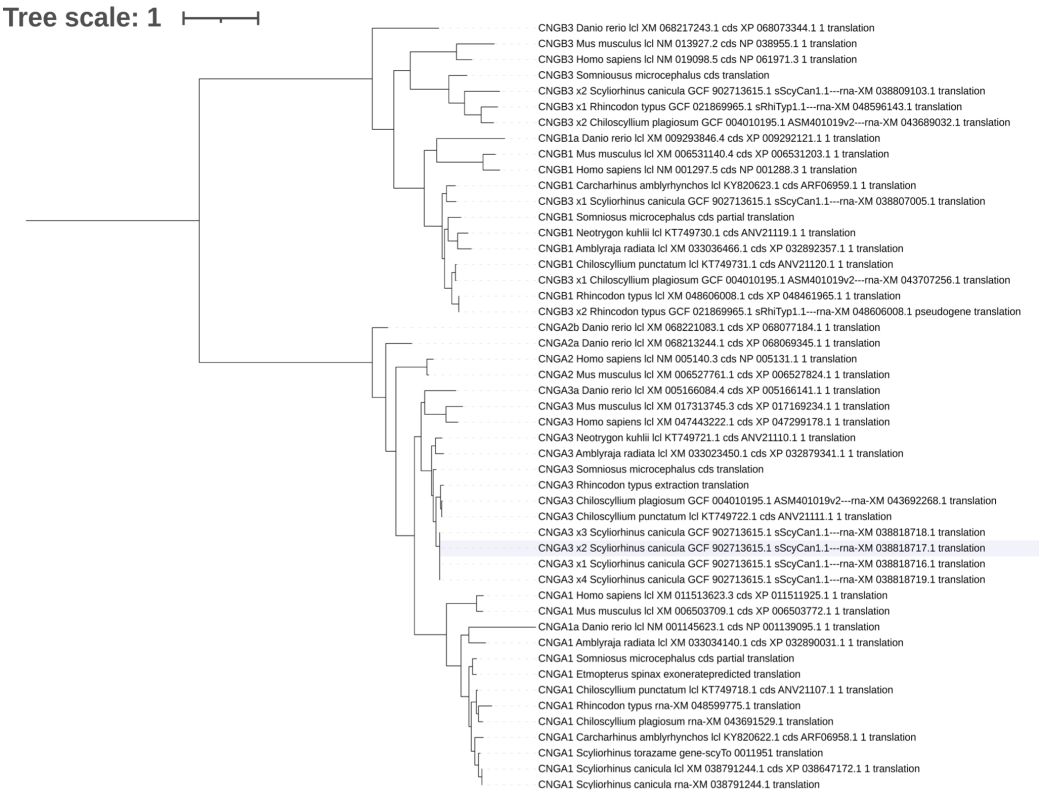


**Fig S3. Cyclic nucleotide gated channel phylogeny.** Gene tree with sequences from several shark species and outgroup species (*Danio rerio, Homo sapiens* and *Mus musculus*). The scale bar denotes substitutions per site.

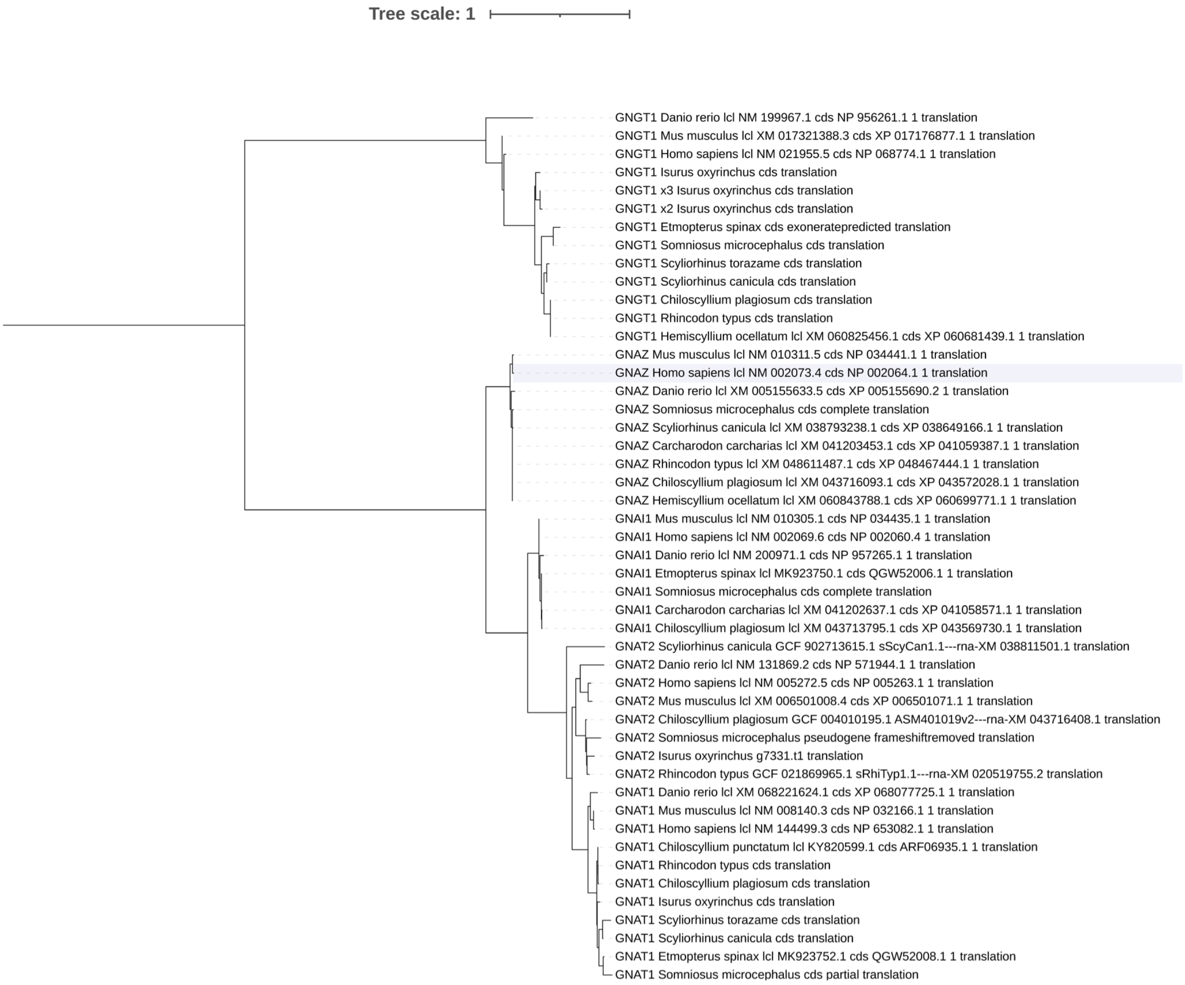


**Fig S4. Transducin phylogeny.** Gene tree with sequences from several shark species and outgroup species (*Danio rerio, Homo sapiens* and *Mus musculus*). The scale bar denotes substitutions per site.

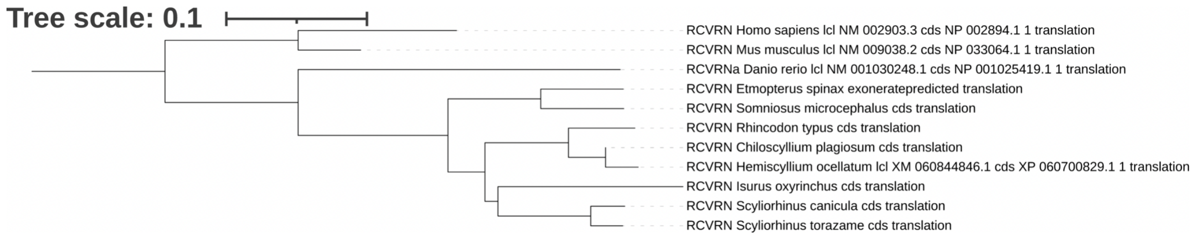


**Fig S5. Recoverin phylogeny.** Gene tree with sequences from several shark species and outgroup species (*Danio rerio, Homo sapiens* and *Mus musculus*). The scale bar denotes substitutions per site.

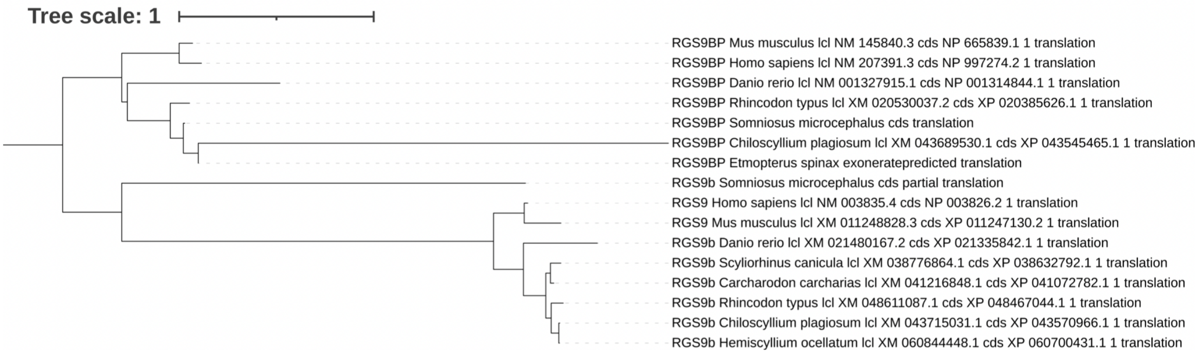


**Fig S6. Regulator of G protein signalling 9 phylogeny.** Gene tree with sequences from several shark species and outgroup species (*Danio rerio, Homo sapiens* and *Mus musculus*). The scale bar denotes substitutions per site.

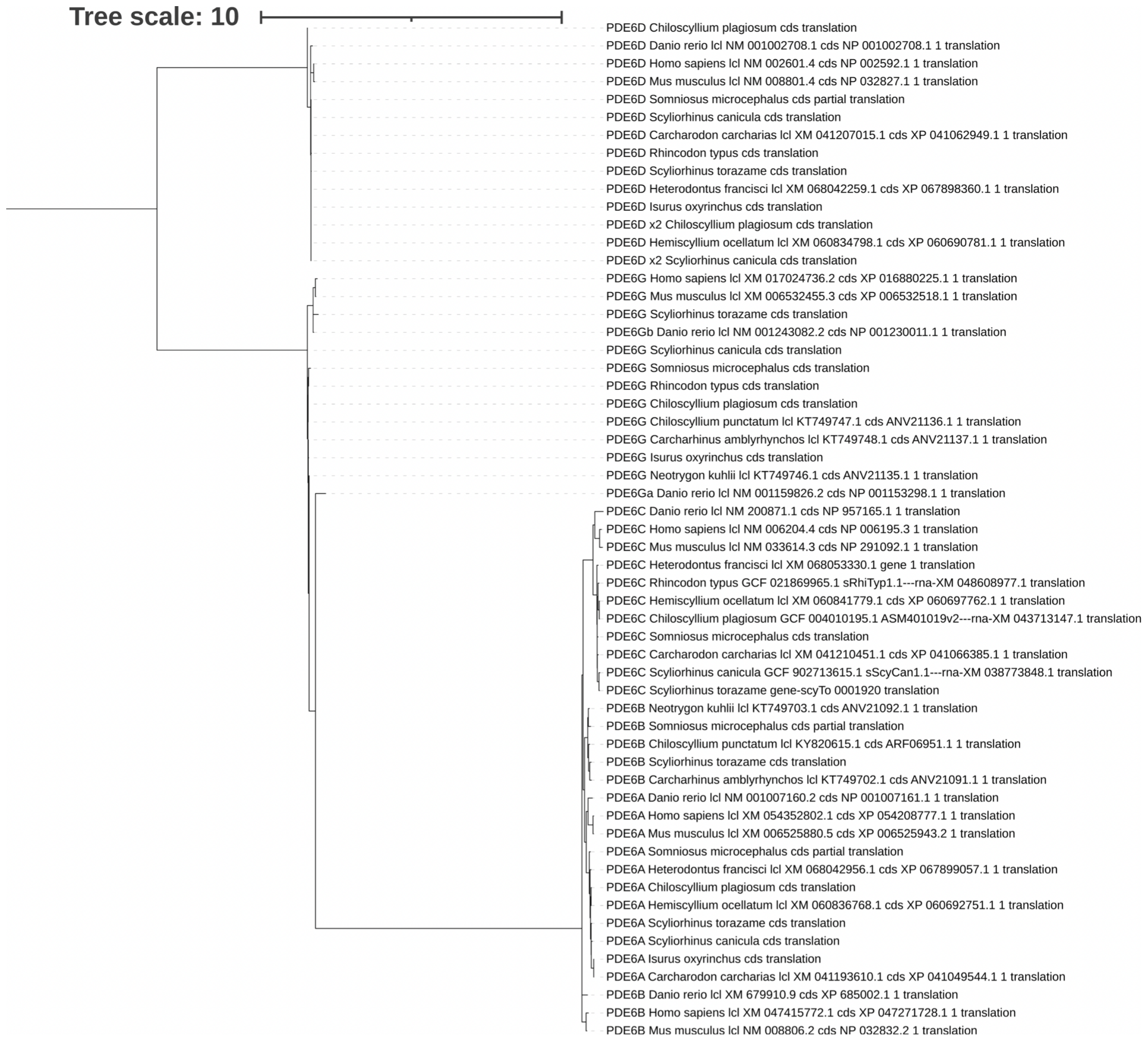


**Fig S7. Phosphodiesterase phylogeny.** Gene tree with sequences from several shark species and outgroup species (*Danio rerio, Homo sapiens* and *Mus musculus*). The scale bar denotes substitutions per site.

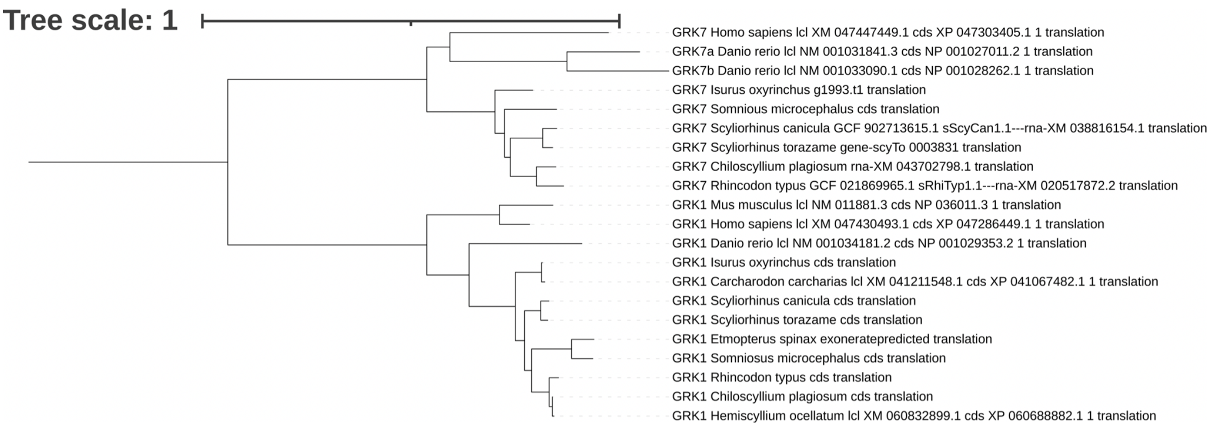


**Fig S8. G-protein coupled receptor kinase phylogeny.** Gene tree with sequences from several shark species and outgroup species (*Danio rerio, Homo sapiens* and *Mus musculus*). The scale bar denotes substitutions per site.

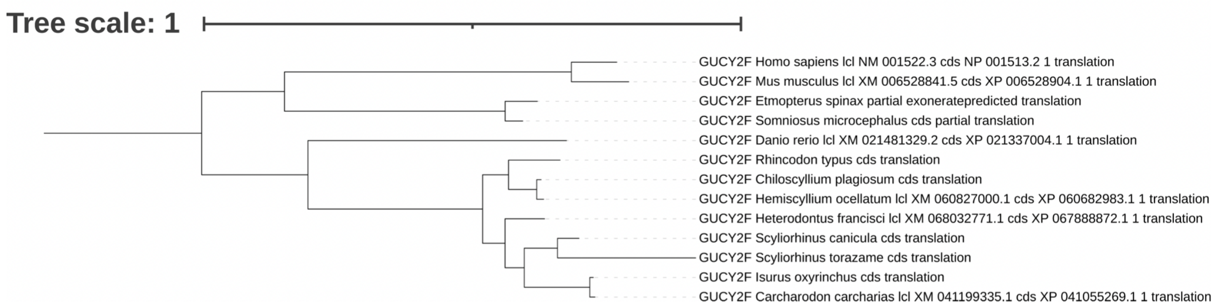


**Fig S9. Guanylate cyclase 2F phylogeny.** Gene tree with sequences from several shark species and outgroup species (*Danio rerio, Homo sapiens* and *Mus musculus*). The scale bar denotes substitutions per site.

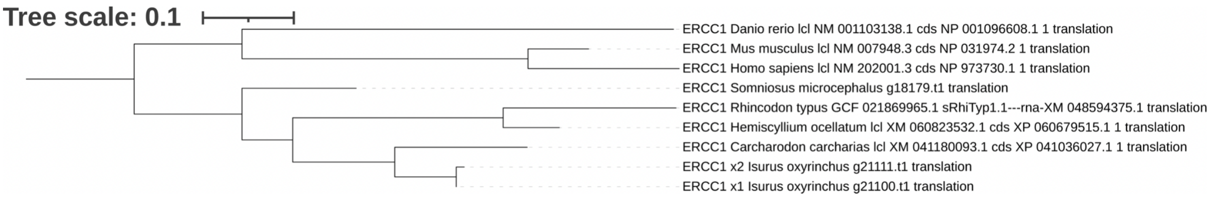


**Fig S10. ERCC excision repair 1 phylogeny.** Gene tree with sequences from several shark species and outgroup species (*Danio rerio, Homo sapiens* and *Mus musculus*). The scale bar denotes substitutions per site.

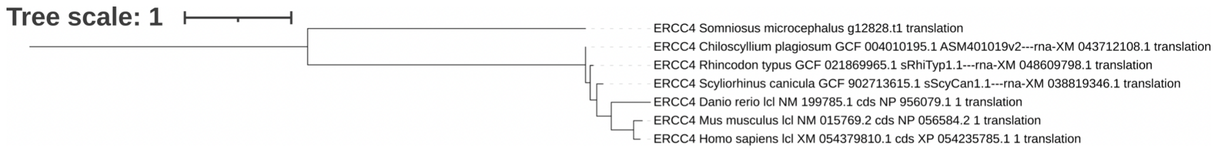


**Fig S11. ERCC excision repair 4 phylogeny.** Gene tree with sequences from several shark species and outgroup species (*Danio rerio, Homo sapiens* and *Mus musculus*). The scale bar denotes substitutions per site.

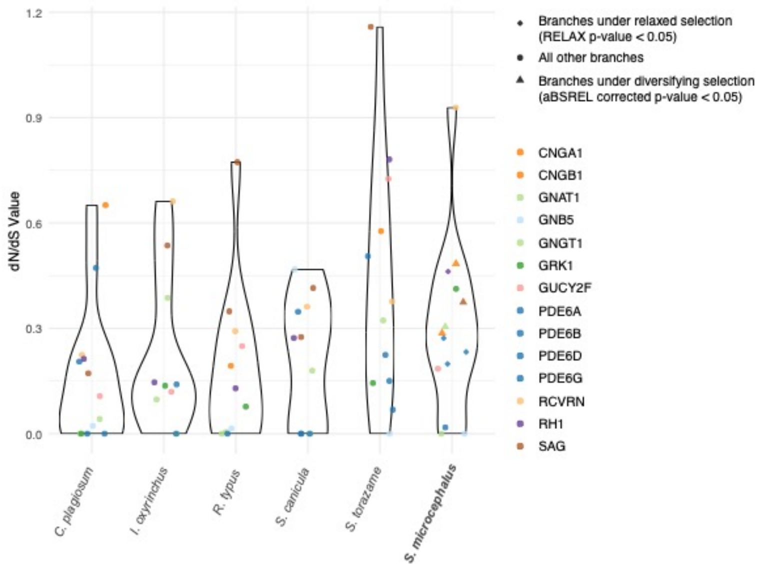


**Fig S12. Distribution of dN/dS (ω) values of rod phototransduction genes in chondrichthyan species.** Violin plots showing per-species distributions of ω ratios for genes involved in rod-based phototransduction, coloured by gene type.

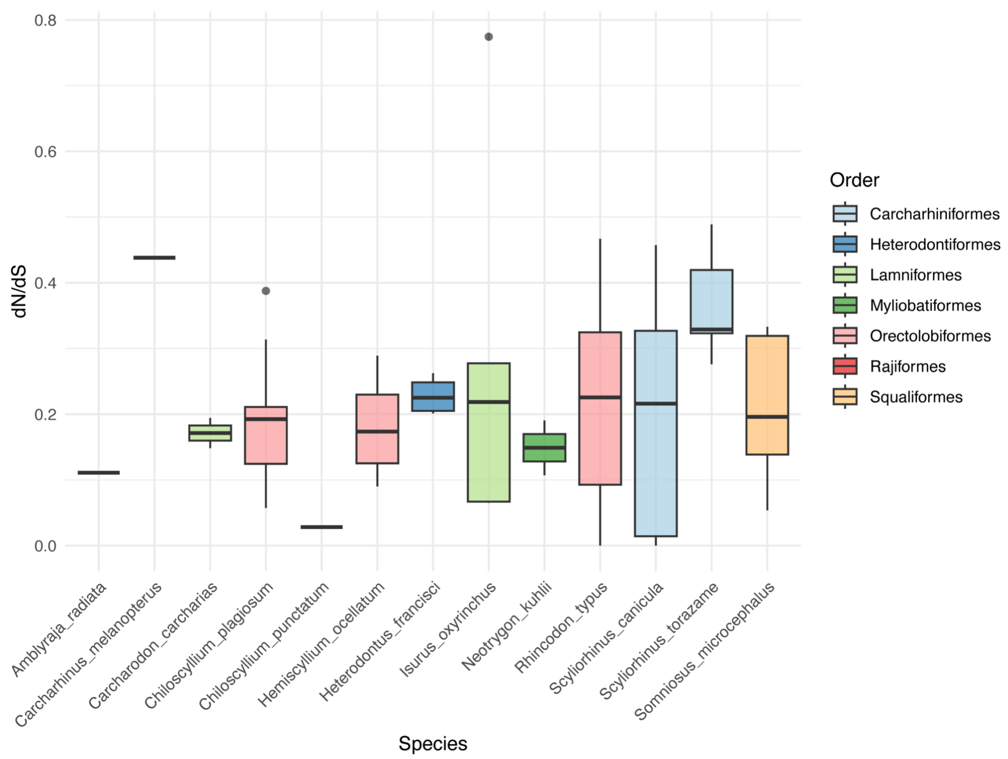


**Fig S13. Distribution of dN/dS (ω) values of all retrieved cone and non-visual phototransduction genes in chondrichthyan species.** Box-and-whisker plots showing per-species distributions of ω ratios for genes involved in cone-based and non-visual phototransduction, coloured by order.

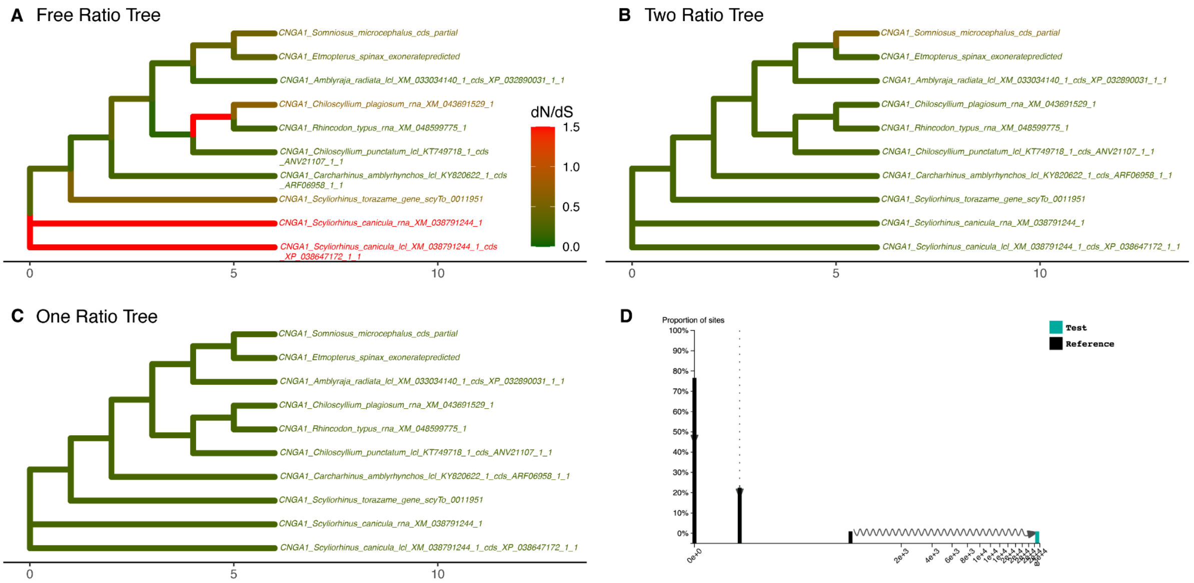


E

| Model | np | lnL |
|---|---|---|
| FreeRatio | 35 | −7129.829 |
| OneRatio | 19 | −7150.629 |
| TwoRatio | 20 | −7147.765 |

F

| Comparison | LRT | df | p.value |
|---|---|---|---|
| TwoRatio vs OneRatio | 5.727874 | 1 | 0.0166976855 |
| FreeRatio vs OneRatio | 41.599588 | 16 | 0.0004526283 |
| FreeRatio vs TwoRatio | 35.871714 | 15 | 0.0018447019 |

**Fig S14. Natural selection analysis on chondrichthyan CNGA1 genes.** PAML was used to compute the dN/dS (= ω) per branch under four different models. **A**, Free-ratio model which fits one ω ratio per branch. **B**, Two-ratio model with one ω ratio for *Somniosus microcephalus* and one ω ratio for other shark species. **C**, One-ratio model with the same ω ratio for all shark species. **D**, RELAX ω distribution results under the RELAX alternative model, assigning *Somniosus microcephalus* as test branches and all other shark species as reference branches. **E**, Log-likelihood (lnL) and number of parameters (np) of each model. **F**, Log-likelihood ratio test (LRT) between each of the models, showing degrees of freedom (df) and statistical significance (p.value) for each comparison.

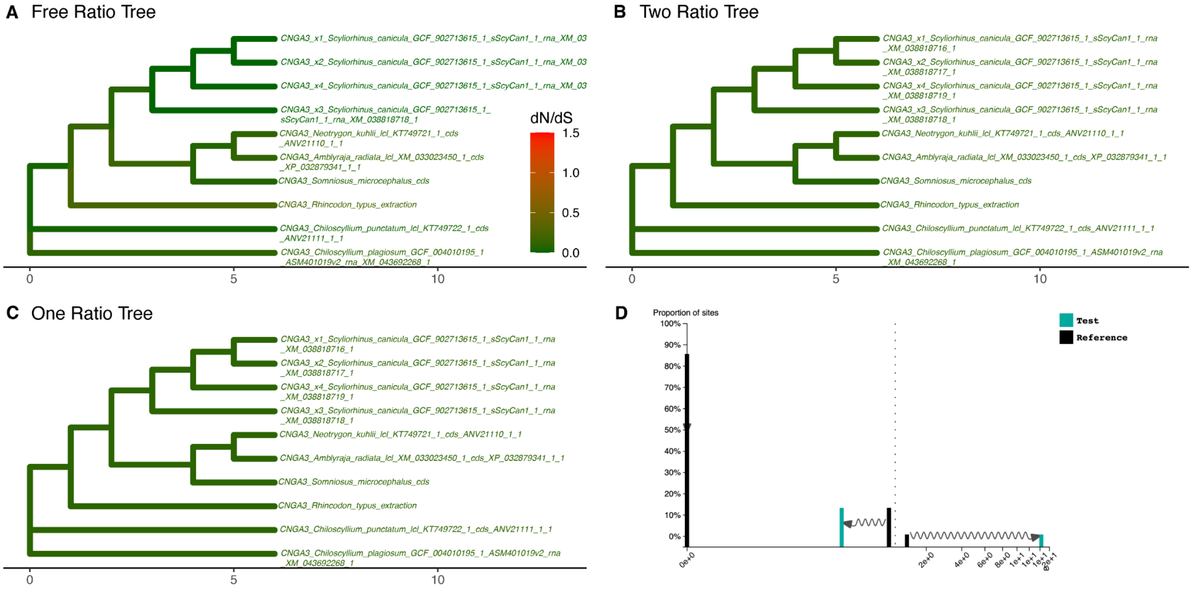


| | Model | np | lnL |
|---|---|---|---|
| E | FreeRatio | 35 | −6059.964 |
| | OneRatio | 19 | −6070.006 |
| | TwoRatio | 20 | −6069.474 |

| | Comparison | LRT | df | p.value |
|---|---|---|---|---|
| F | TwoRatio vs OneRatio | 1.06385 | 1 | 0.3023387 |
| | FreeRatio vs OneRatio | 20.08317 | 16 | 0.2164979 |
| | FreeRatio vs TwoRatio | 19.01932 | 15 | 0.2128595 |

**Fig S15. Natural selection analysis on chondrichthyan CNGA3 genes.** PAML was used to compute the dN/dS (= ω) per branch under four different models. **A**, Free-ratio model which fits one ω ratio per branch. **B**, Two-ratio model with one ω ratio for *Somniosus microcephalus* and one ω ratio for other shark species. **C**, One-ratio model with the same ω ratio for all shark species. **D**, RELAX ω distribution results under the RELAX alternative model, assigning *Somniosus microcephalus* as test branches and all other shark species as reference branches. **E**, Log-likelihood (lnL) and number of parameters (np) of each model. **F**, Log-likelihood ratio test (LRT) between each of the models, showing degrees of freedom (df) and statistical significance (p.value) for each comparison.

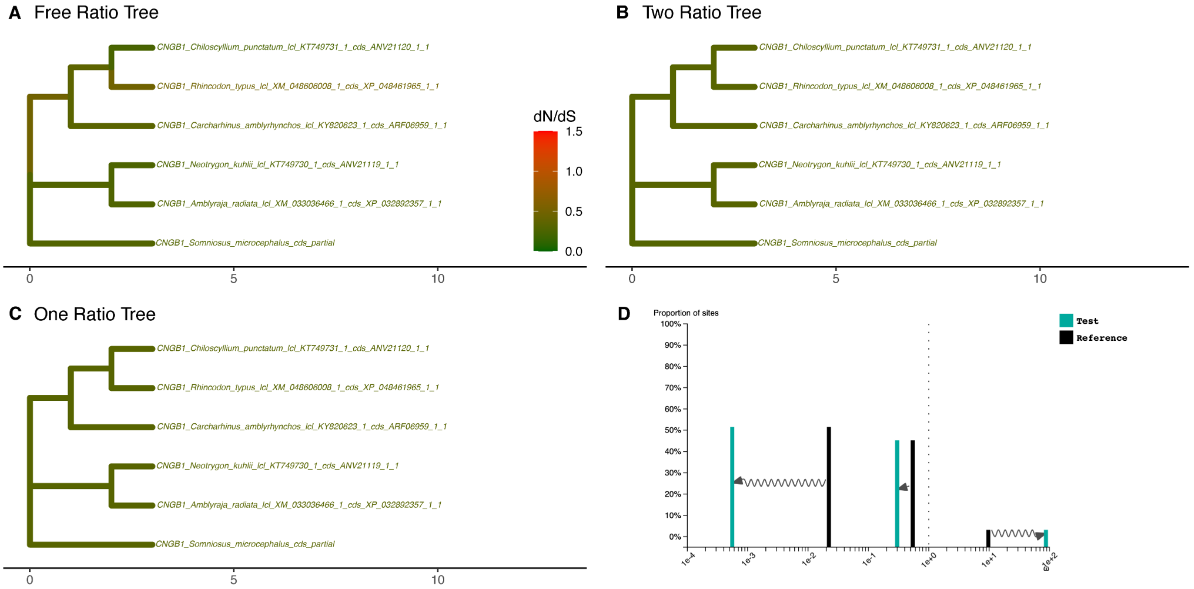


| Model | np | lnL |
|---|---|---|
| FreeRatio | 19 | −14210.24 |
| OneRatio | 11 | −14215.58 |
| TwoRatio | 12 | −14215.55 |

| Comparison | LRT | df | p.value |
|---|---|---|---|
| TwoRatio vs OneRatio | 0.063506 | 1 | 0.8010380 |
| FreeRatio vs OneRatio | 10.671648 | 8 | 0.2210070 |
| FreeRatio vs TwoRatio | 10.608142 | 7 | 0.1566485 |

**Fig S16. Natural selection analysis on chondrichthyan CNGB1 genes.** PAML was used to compute the dN/dS (= ω) per branch under four different models. **A**, Free-ratio model which fits one ω ratio per branch. **B**, Two-ratio model with one ω ratio for *Somniosus microcephalus* and one ω ratio for other shark species. **C**, One-ratio model with the same ω ratio for all shark species. **D**, RELAX ω distribution results under the RELAX alternative model, assigning *Somniosus microcephalus* as test branches and all other shark species as reference branches. **E**, Log-likelihood (lnL) and number of parameters (np) of each model. **F**, Log-likelihood ratio test (LRT) between each of the models, showing degrees of freedom (df) and statistical significance (p.value) for each comparison.

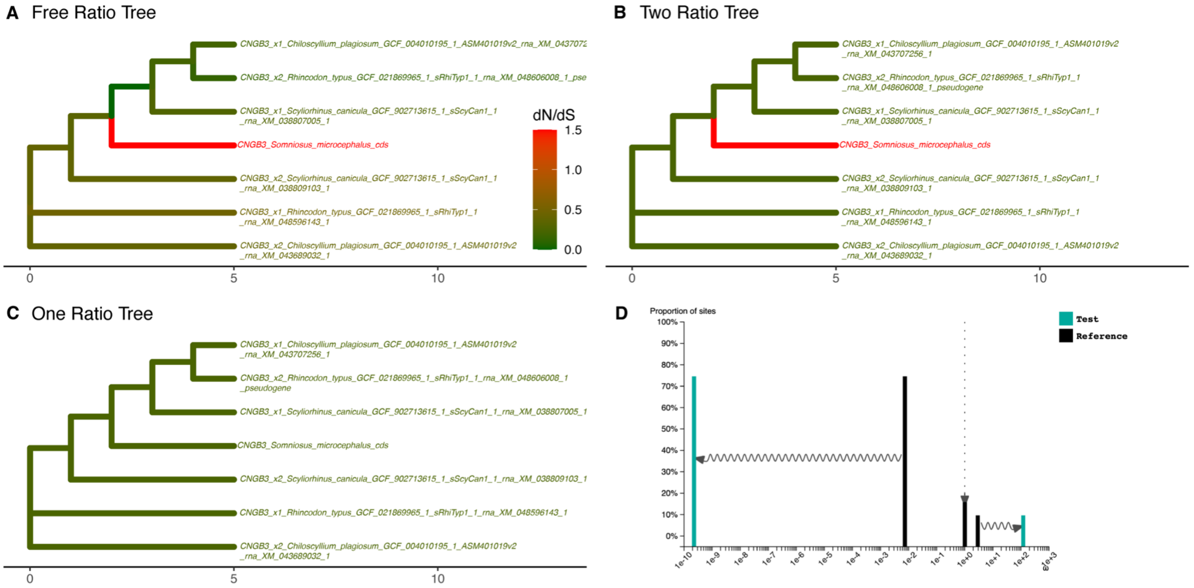


E

| Model | np | lnL |
|---|---|---|
| FreeRatio | 23 | −10619.86 |
| OneRatio | 13 | −10677.26 |
| TwoRatio | 14 | −10674.20 |

F

| Comparison | LRT | df | p.value |
|---|---|---|---|
| TwoRatio vs OneRatio | 6.111974 | 1 | 1.342691e−02 |
| FreeRatio vs OneRatio | 114.791764 | 10 | 5.746244e−20 |
| FreeRatio vs TwoRatio | 108.679790 | 9 | 2.729635e−19 |

**Fig S17. Natural selection analysis on chondrichthyan CNGB3 genes.** PAML was used to compute the dN/dS (= ω) per branch under four different models. **A**, Free-ratio model which fits one ω ratio per branch. **B**, Two-ratio model with one ω ratio for *Somniosus microcephalus* and one ω ratio for other shark species. **C**, One-ratio model with the same ω ratio for all shark species. **D**, RELAX ω distribution results under the RELAX alternative model, assigning *Somniosus microcephalus* as test branches and all other shark species as reference branches. **E**, Log-likelihood (lnL) and number of parameters (np) of each model. **F**, Log-likelihood ratio test (LRT) between each of the models, showing degrees of freedom (df) and statistical significance (p.value) for each comparison.

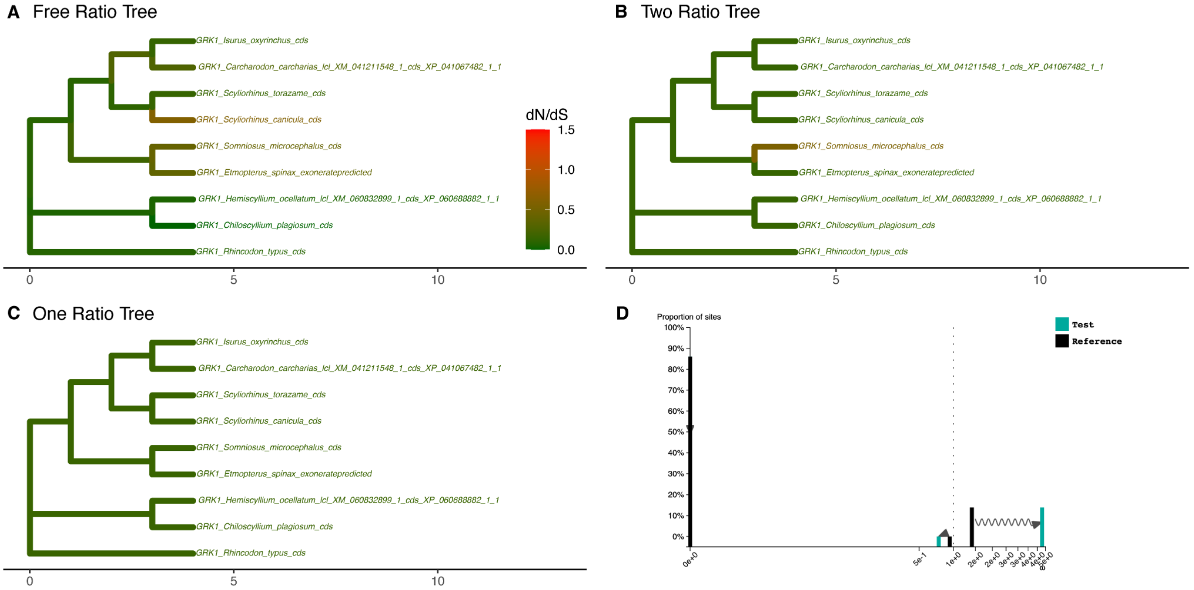


**E**

| Model | np | lnL |
|---|---|---|
| FreeRatio | 31 | −5321.285 |
| OneRatio | 17 | −5348.864 |
| TwoRatio | 18 | −5341.773 |

**F**

| Comparison | LRT | df | p.value |
|---|---|---|---|
| TwoRatio vs OneRatio | 14.18285 | 1 | 1.658753e−04 |
| FreeRatio vs OneRatio | 55.15839 | 14 | 8.132058e−07 |
| FreeRatio vs TwoRatio | 40.97554 | 13 | 9.616008e−05 |

**Fig S18. Natural selection analysis on chondrichthyan GRK1 genes.** PAML was used to compute the dN/dS (= ω) per branch under four different models. **A**, Free-ratio model which fits one ω ratio per branch. **B**, Two-ratio model with one ω ratio for *Somniosus microcephalus* and one ω ratio for other shark species. **C**, One-ratio model with the same ω ratio for all shark species. **D**, RELAX ω distribution results under the RELAX alternative model, assigning *Somniosus microcephalus* as test branches and all other shark species as reference branches. **E**, Log-likelihood (lnL) and number of parameters (np) of each model. **F**, Log-likelihood ratio test (LRT) between each of the models, showing degrees of freedom (df) and statistical significance (p.value) for each comparison.

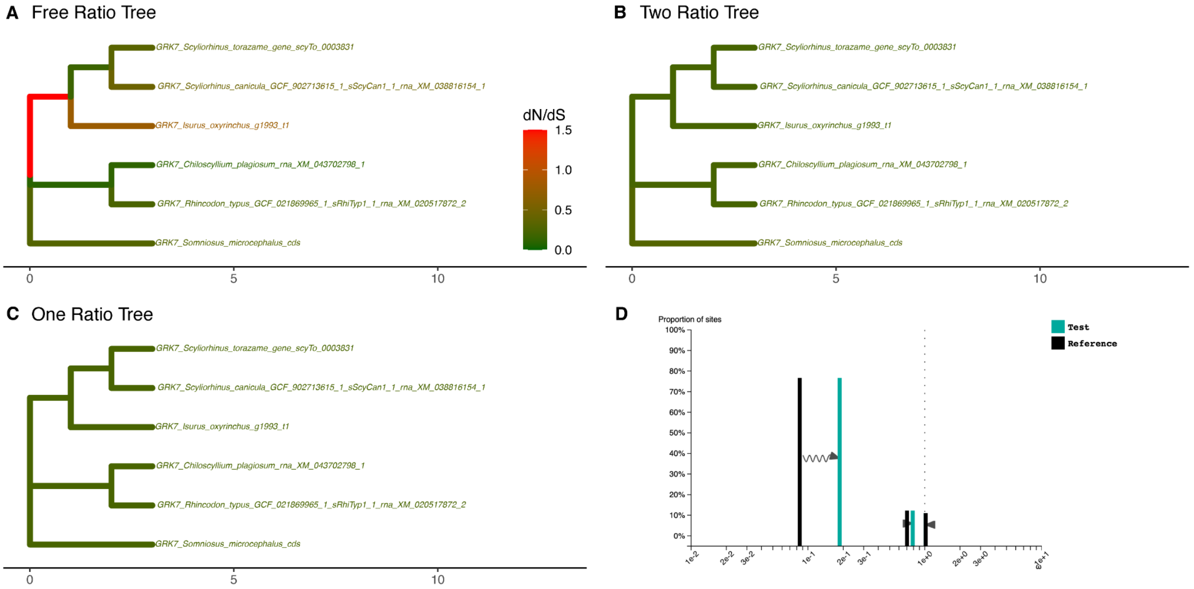


E

| Model | np | lnL |
|---|---|---|
| FreeRatio | 19 | −4725.883 |
| OneRatio | 11 | −4749.558 |
| TwoRatio | 12 | −4749.034 |

F

| Comparison | LRT | df | p.value |
|---|---|---|---|
| TwoRatio vs OneRatio | 1.047236 | 1 | 3.061445e−01 |
| FreeRatio vs OneRatio | 47.348784 | 8 | 1.315596e−07 |
| FreeRatio vs TwoRatio | 46.301548 | 7 | 7.636787e−08 |

**Fig S19. Natural selection analysis on chondrichthyan GRK7 genes.** PAML was used to compute the dN/dS (= ω) per branch under four different models. **A**, Free-ratio model which fits one ω ratio per branch. **B**, Two-ratio model with one ω ratio for *Somniosus microcephalus* and one ω ratio for other shark species. **C**, One-ratio model with the same ω ratio for all shark species. **D**, RELAX ω distribution results under the RELAX alternative model, assigning *Somniosus microcephalus* as test branches and all other shark species as reference branches. **E**, Log-likelihood (lnL) and number of parameters (np) of each model. **F**, Log-likelihood ratio test (LRT) between each of the models, showing degrees of freedom (df) and statistical significance (p.value) for each comparison.

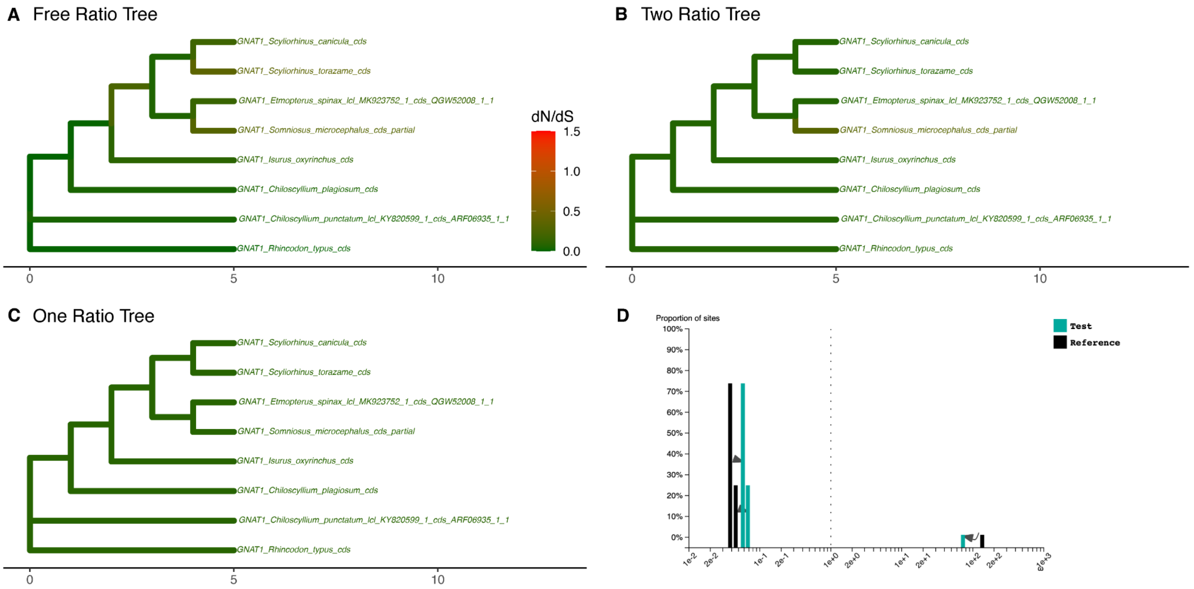


| E | Model | np | lnL |
|---|---|---|---|
| | FreeRatio | 27 | −2944.274 |
| | OneRatio | 15 | −2986.261 |
| | TwoRatio | 16 | −2977.586 |

| F | Comparison | LRT | df | p.value |
|---|---|---|---|---|
| | TwoRatio vs OneRatio | 17.34884 | 1 | 3.110874e−05 |
| | FreeRatio vs OneRatio | 83.97279 | 12 | 7.169221e−13 |
| | FreeRatio vs TwoRatio | 66.62395 | 11 | 5.327022e−10 |

**Fig S20. Natural selection analysis on chondrichthyan GNAT1 genes.** PAML was used to compute the dN/dS (= ω) per branch under four different models. **A**, Free-ratio model which fits one ω ratio per branch. **B**, Two-ratio model with one ω ratio for *Somniosus microcephalus* and one ω ratio for other shark species. **C**, One-ratio model with the same ω ratio for all shark species. **D**, RELAX ω distribution results under the RELAX alternative model, assigning *Somniosus microcephalus* as test branches and all other shark species as reference branches. **E**, Log-likelihood (lnL) and number of parameters (np) of each model. **F**, Log-likelihood ratio test (LRT) between each of the models, showing degrees of freedom (df) and statistical significance (p.value) for each comparison.

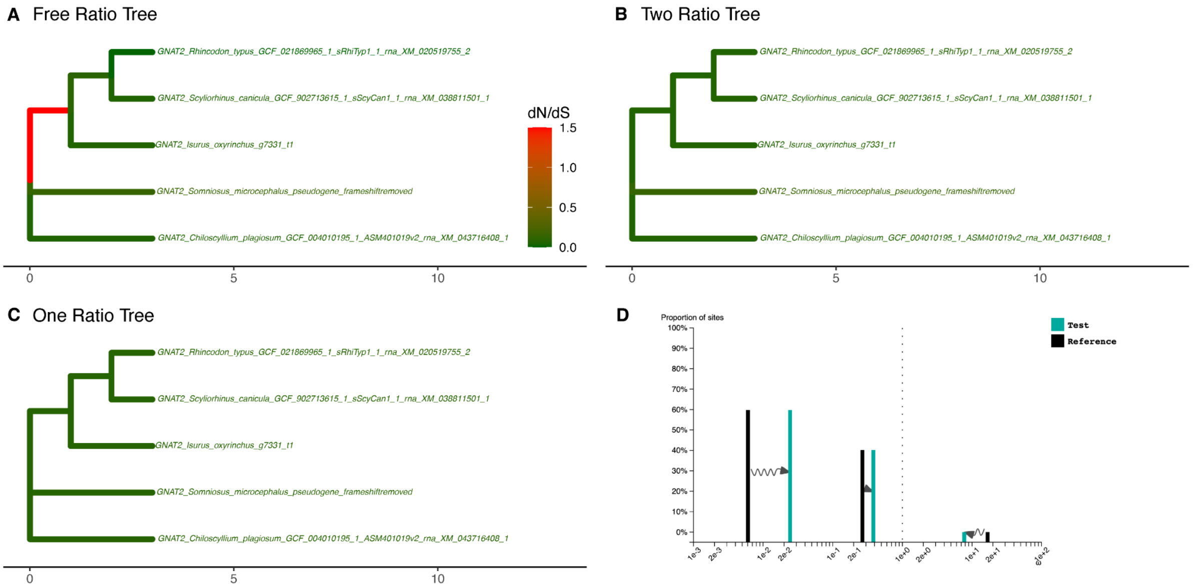


| E | Model | np | lnL |
|---|---|---|---|
| | FreeRatio | 15 | –3058.983 |
| | OneRatio | 9 | –3062.905 |
| | TwoRatio | 10 | –3059.995 |

| F | Comparison | LRT | df | p.value |
|---|---|---|---|---|
| | TwoRatio vs OneRatio | 5.820942 | 1 | 0.01583646 |
| | FreeRatio vs OneRatio | 7.843810 | 6 | 0.24977102 |
| | FreeRatio vs TwoRatio | 2.022868 | 5 | 0.84597185 |

**Fig S21. Natural selection analysis on chondrichthyan GNAT2 genes.** PAML was used to compute the dN/dS (= ω) per branch under four different models. **A**, Free-ratio model which fits one ω ratio per branch. **B**, Two-ratio model with one ω ratio for *Somniosus microcephalus* and one ω ratio for other shark species. **C**, One-ratio model with the same ω ratio for all shark species. **D**, RELAX ω distribution results under the RELAX alternative model, assigning *Somniosus microcephalus* as test branches and all other shark species as reference branches. **E**, Log-likelihood (lnL) and number of parameters (np) of each model. **F**, Log-likelihood ratio test (LRT) between each of the models, showing degrees of freedom (df) and statistical significance (p.value) for each comparison.

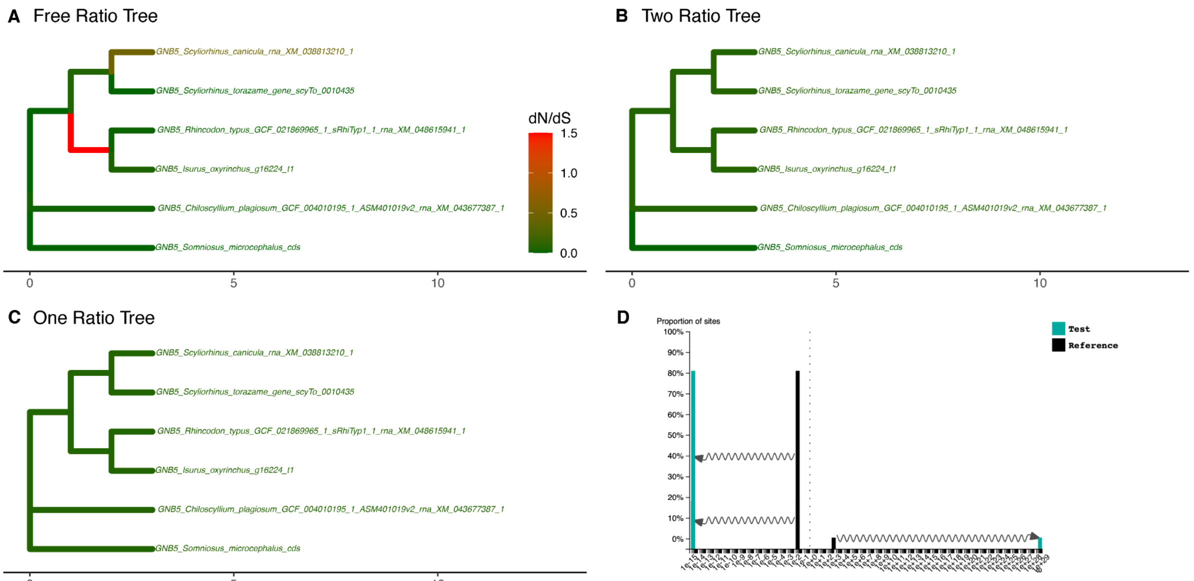


E

| Model | np | lnL |
|---|---|---|
| FreeRatio | 19 | −2527.125 |
| OneRatio | 11 | −2550.132 |
| TwoRatio | 12 | −2548.493 |

F

| Comparison | LRT | df | p.value |
|---|---|---|---|
| TwoRatio vs OneRatio | 3.277078 | 1 | 7.025390e−02 |
| FreeRatio vs OneRatio | 46.014280 | 8 | 2.362182e−07 |
| FreeRatio vs TwoRatio | 42.737202 | 7 | 3.748896e−07 |

**Fig S22. Natural selection analysis on chondrichthyan GNB5 genes.** PAML was used to compute the dN/dS (= ω) per branch under four different models. **A**, Free-ratio model which fits one ω ratio per branch. **B**, Two-ratio model with one ω ratio for *Somniosus microcephalus* and one ω ratio for other shark species. **C**, One-ratio model with the same ω ratio for all shark species. **D**, RELAX ω distribution results under the RELAX alternative model, assigning *Somniosus microcephalus* as test branches and all other shark species as reference branches. **E**, Log-likelihood (lnL) and number of parameters (np) of each model. **F**, Log-likelihood ratio test (LRT) between each of the models, showing degrees of freedom (df) and statistical significance (p.value) for each comparison.

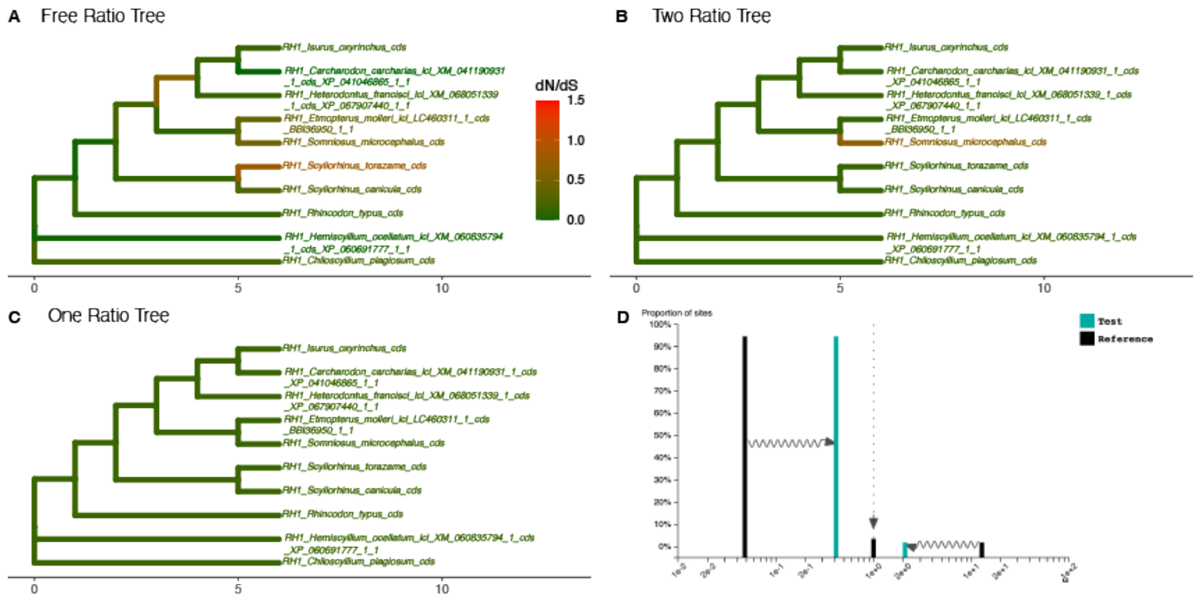


E

| Model | np | lnL |
|---|---|---|
| FreeRatio | 35 | −3960.812 |
| OneRatio | 19 | −3998.088 |
| TwoRatio | 20 | −3991.196 |

F

| Comparison | LRT | df | p.value |
|---|---|---|---|
| TwoRatio vs OneRatio | 13.78547 | 1 | 2.049151e−04 |
| FreeRatio vs OneRatio | 74.55243 | 16 | 1.570365e−09 |
| FreeRatio vs TwoRatio | 60.76696 | 15 | 1.860789e−07 |

**Fig S23. Natural selection analysis on chondrichthyan RH1 genes.** PAML was used to compute the dN/dS (= ω) per branch under four different models. **A**, Free-ratio model which fits one ω ratio per branch. **B**, Two-ratio model with one ω ratio for *Somniosus microcephalus* and one ω ratio for other shark species. **C**, One-ratio model with the same ω ratio for all shark species. **D**, RELAX ω distribution results under the RELAX alternative model, assigning *Somniosus microcephalus* as test branches and all other shark species as reference branches. **E**, Log-likelihood (lnL) and number of parameters (np) of each model. **F**, Log-likelihood ratio test (LRT) between each of the models, showing degrees of freedom (df) and statistical significance (p.value) for each comparison.

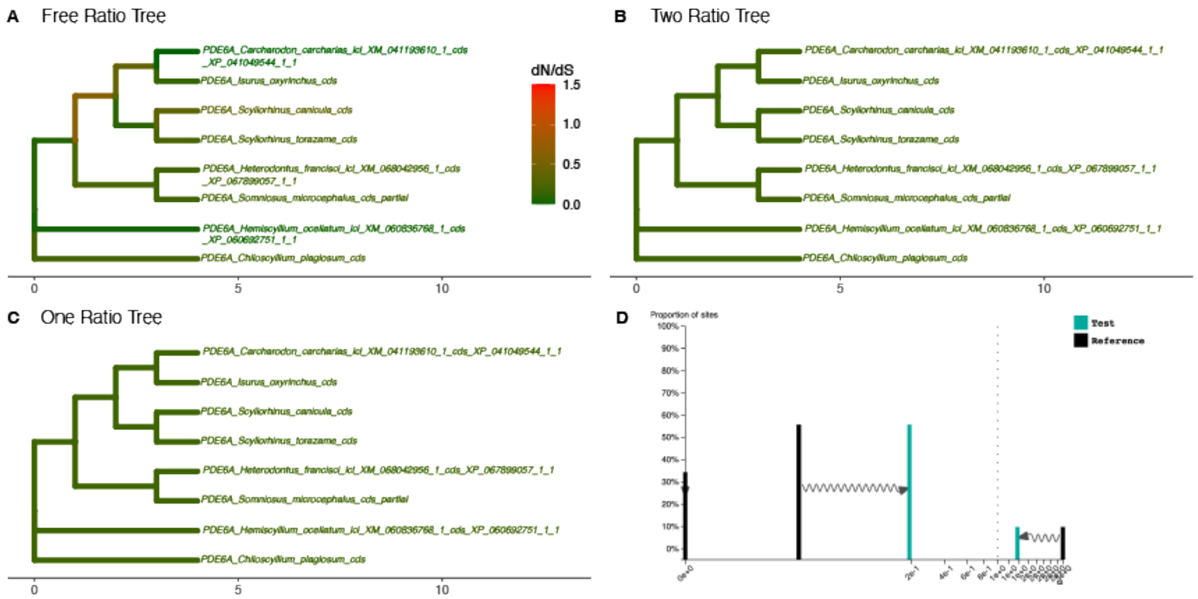


E

| Model | np | lnL |
|---|---|---|
| FreeRatio | 27 | −6635.944 |
| OneRatio | 15 | −6665.061 |
| TwoRatio | 16 | −6664.966 |

F

| Comparison | LRT | df | p.value |
|---|---|---|---|
| TwoRatio vs OneRatio | 0.18989 | 1 | 6.630082e−01 |
| FreeRatio vs OneRatio | 58.23436 | 12 | 4.726730e−08 |
| FreeRatio vs TwoRatio | 58.04447 | 11 | 2.135206e−08 |

**Fig S24. Natural selection analysis on chondrichthyan PDE6A genes.** PAML was used to compute the dN/dS (= ω) per branch under four different models. **A**, Free-ratio model which fits one ω ratio per branch. **B**, Two-ratio model with one ω ratio for *Somniosus microcephalus* and one ω ratio for other shark species. **C**, One-ratio model with the same ω ratio for all shark species. **D**, RELAX ω distribution results under the RELAX alternative model, assigning *Somniosus microcephalus* as test branches and all other shark species as reference branches. **E**, Log-likelihood (lnL) and number of parameters (np) of each model. **F**, Log-likelihood ratio test (LRT) between each of the models, showing degrees of freedom (df) and statistical significance (p.value) for each comparison.

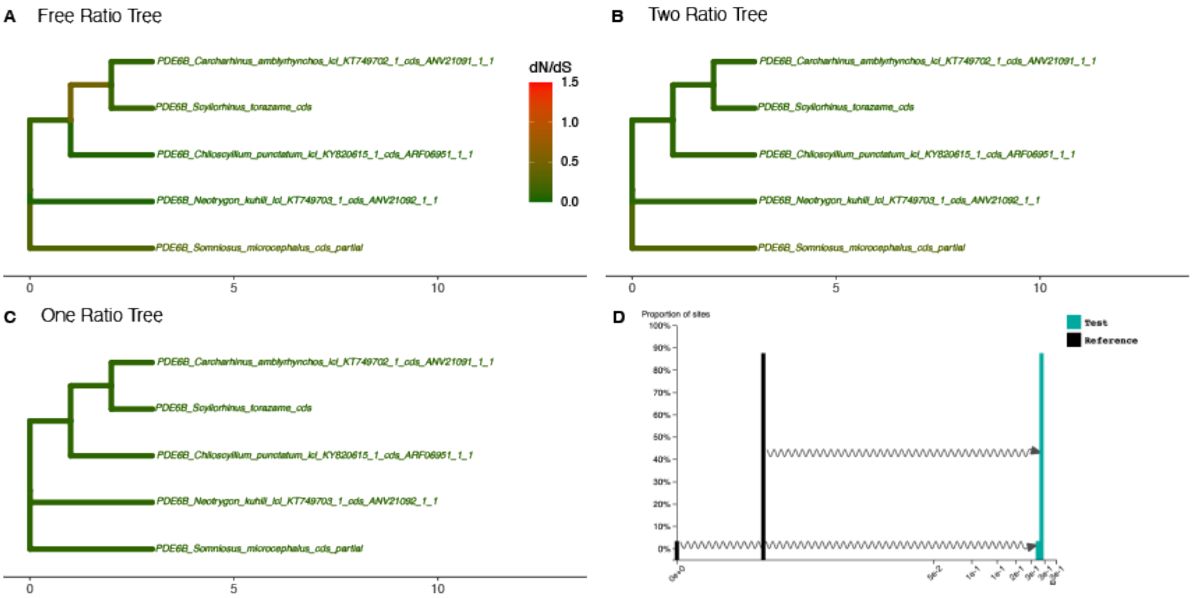


E

| Model | np | lnL |
|---|---|---|
| FreeRatio | 15 | −6489.430 |
| OneRatio | 9 | −6506.985 |
| TwoRatio | 10 | −6502.790 |

F

| Comparison | LRT | df | p.value |
|---|---|---|---|
| TwoRatio vs OneRatio | 8.39019 | 1 | 3.772515e−03 |
| FreeRatio vs OneRatio | 35.11175 | 6 | 4.099891e−06 |
| FreeRatio vs TwoRatio | 26.72156 | 5 | 6.462342e−05 |

**Fig S25. Natural selection analysis on chondrichthyan PDE6B genes.** PAML was used to compute the dN/dS (= ω) per branch under four different models. **A**, Free-ratio model which fits one ω ratio per branch. **B**, Two-ratio model with one ω ratio for *Somniosus microcephalus* and one ω ratio for other shark species. **C**, One-ratio model with the same ω ratio for all shark species. **D**, RELAX ω distribution results under the RELAX alternative model, assigning *Somniosus microcephalus* as test branches and all other shark species as reference branches. **E**, Log-likelihood (lnL) and number of parameters (np) of each model. **F**, Log-likelihood ratio test (LRT) between each of the models, showing degrees of freedom (df) and statistical significance (p.value) for each comparison.

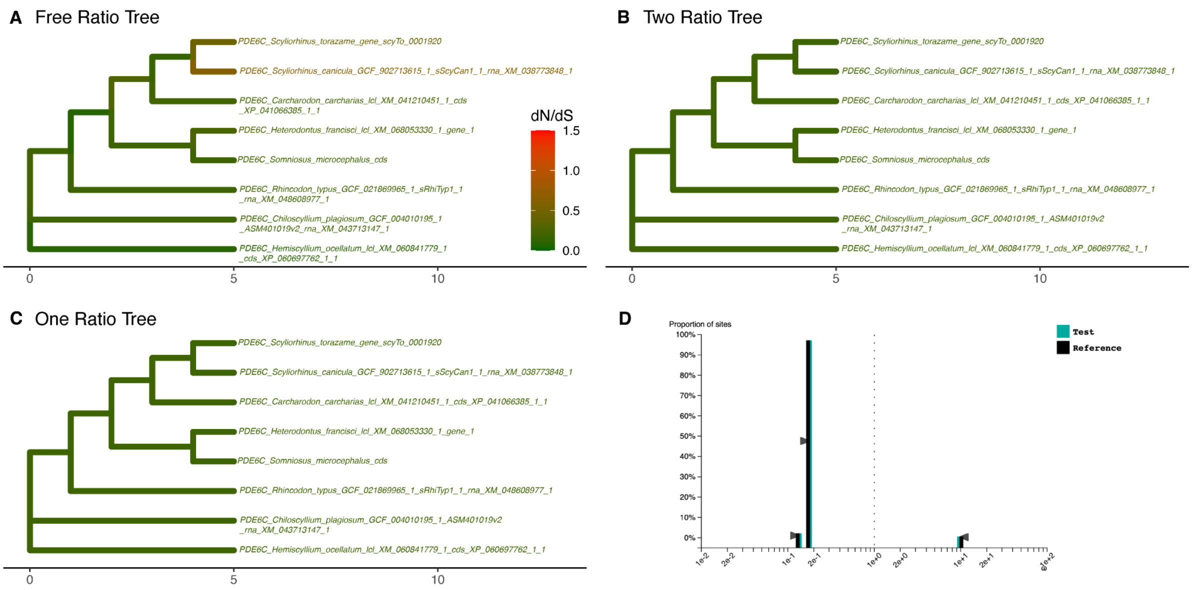


E

| Model | np | lnL |
|---|---|---|
| FreeRatio | 27 | –6761.486 |
| OneRatio | 15 | –6774.442 |
| TwoRatio | 16 | –6774.438 |

F

| Comparison | LRT | df | p.value |
|---|---|---|---|
| TwoRatio vs OneRatio | 0.008598 | 1 | 0.926121717 |
| FreeRatio vs OneRatio | 25.913398 | 12 | 0.011040792 |
| FreeRatio vs TwoRatio | 25.904800 | 11 | 0.006704917 |

**Fig S26. Natural selection analysis on chondrichthyan PDE6C genes.** PAML was used to compute the dN/dS (= ω) per branch under four different models. **A**, Free-ratio model which fits one ω ratio per branch. **B**, Two-ratio model with one ω ratio for *Somniosus microcephalus* and one ω ratio for other shark species. **C**, One-ratio model with the same ω ratio for all shark species. **D**, RELAX ω distribution results under the RELAX alternative model, assigning *Somniosus microcephalus* as test branches and all other shark species as reference branches. **E**, Log-likelihood (lnL) and number of parameters (np) of each model. **F**, Log-likelihood ratio test (LRT) between each of the models, showing degrees of freedom (df) and statistical significance (p.value) for each comparison.

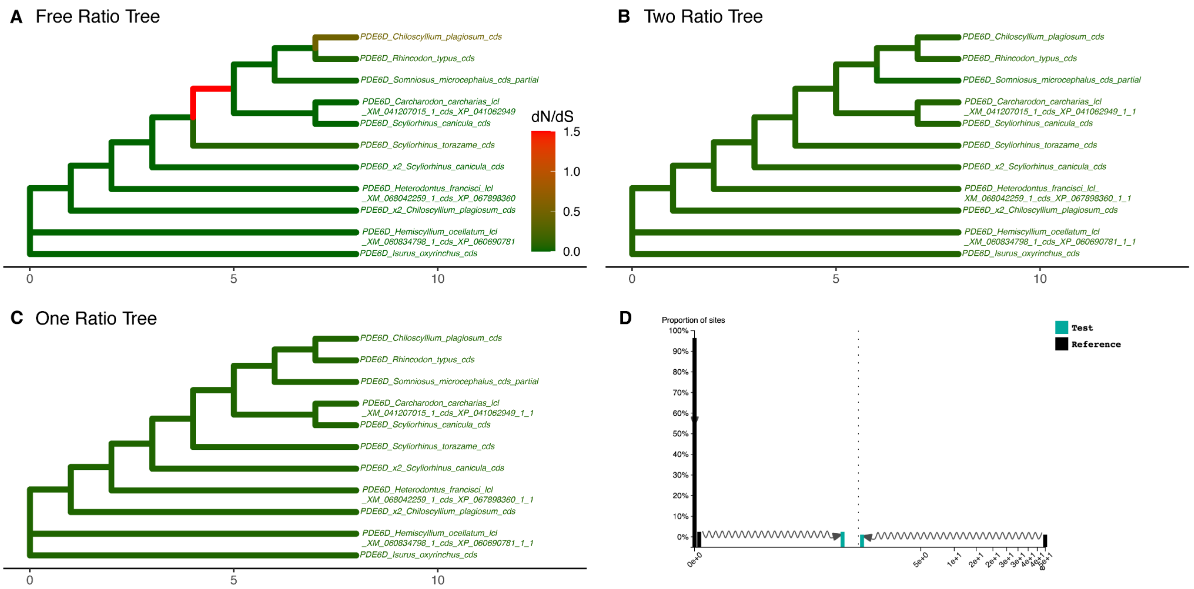


| E | Model | np | lnL |
|---|---|---|---|
| | FreeRatio | 39 | −1257.830 |
| | OneRatio | 21 | −1296.878 |
| | TwoRatio | 22 | −1292.377 |

| F | Comparison | LRT | df | p.value |
|---|---|---|---|---|
| | TwoRatio vs OneRatio | 9.001858 | 1 | 2.697053e−03 |
| | FreeRatio vs OneRatio | 78.096334 | 18 | 1.841249e−09 |
| | FreeRatio vs TwoRatio | 69.094476 | 17 | 3.086404e−08 |

**Fig S27. Natural selection analysis on chondrichthyan PDE6D genes.** PAML was used to compute the dN/dS (= ω) per branch under four different models. **A**, Free-ratio model which fits one ω ratio per branch. **B**, Two-ratio model with one ω ratio for *Somniosus microcephalus* and one ω ratio for other shark species. **C**, One-ratio model with the same ω ratio for all shark species. **D**, RELAX ω distribution results under the RELAX alternative model, assigning *Somniosus microcephalus* as test branches and all other shark species as reference branches. **E**, Log-likelihood (lnL) and number of parameters (np) of each model. **F**, Log-likelihood ratio test (LRT) between each of the models, showing degrees of freedom (df) and statistical significance (p.value) for each comparison.

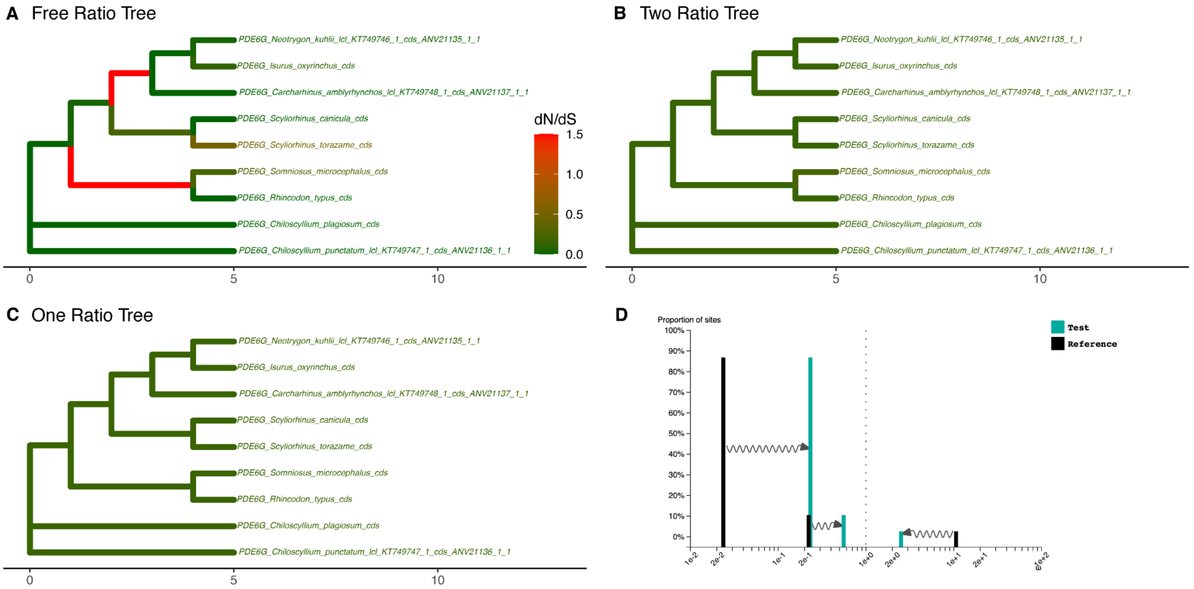


**E**

| Model | np | lnL |
|---|---|---|
| FreeRatio | 31 | –922.9456 |
| OneRatio | 17 | –940.6014 |
| TwoRatio | 18 | –940.3303 |

**F**

| Comparison | LRT | df | p.value |
|---|---|---|---|
| TwoRatio vs OneRatio | 0.54207 | 1 | 0.4615761152 |
| FreeRatio vs OneRatio | 35.31158 | 14 | 0.0013216989 |
| FreeRatio vs TwoRatio | 34.76951 | 13 | 0.0009184926 |

**Fig S28. Natural selection analysis on chondrichthyan PDE6G genes.** PAML was used to compute the dN/dS (= ω) per branch under four different models. **A**, Free-ratio model which fits one ω ratio per branch. **B**, Two-ratio model with one ω ratio for *Somniosus microcephalus* and one ω ratio for other shark species. **C**, One-ratio model with the same ω ratio for all shark species. **D**, RELAX ω distribution results under the RELAX alternative model, assigning *Somniosus microcephalus* as test branches and all other shark species as reference branches. **E**, Log-likelihood (lnL) and number of parameters (np) of each model. **F**, Log-likelihood ratio test (LRT) between each of the models, showing degrees of freedom (df) and statistical significance (p.value) for each comparison.

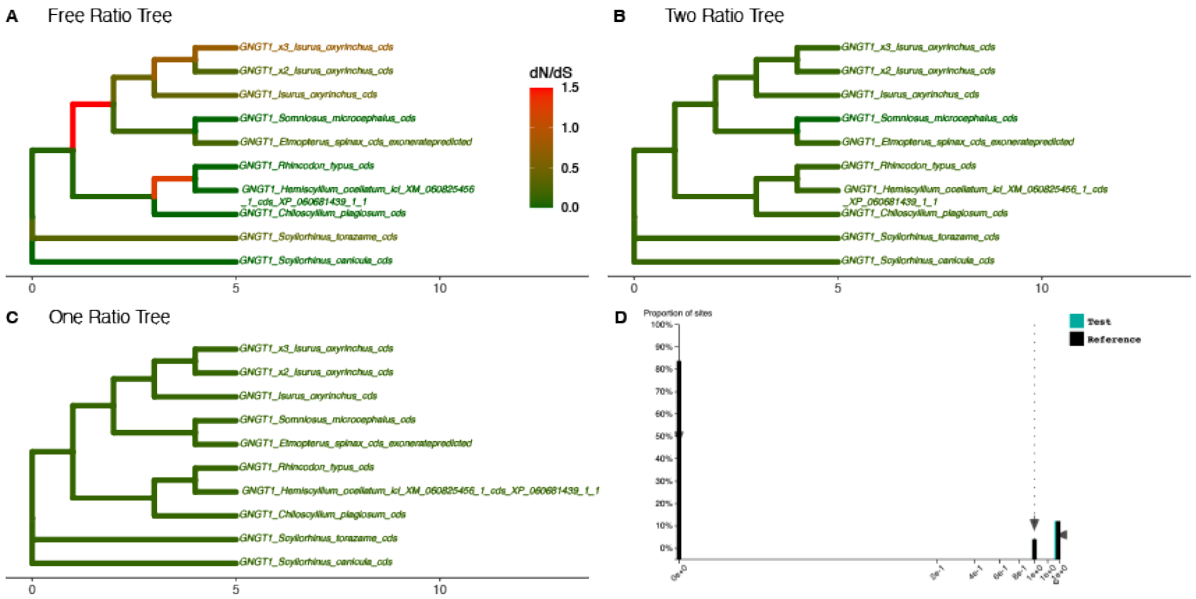


| E | | |
|---|---|---|
| **Model** | **np** | **lnL** |
| FreeRatio | 35 | –579.4468 |
| OneRatio | 19 | –587.9479 |
| TwoRatio | 20 | –587.9479 |

| F | | | |
|---|---|---|---|
| **Comparison** | **LRT** | **df** | **p.value** |
| TwoRatio vs OneRatio | 0.000042 | 1 | 0.9948292 |
| FreeRatio vs OneRatio | 17.002122 | 16 | 0.3854598 |
| FreeRatio vs TwoRatio | 17.002080 | 15 | 0.3187401 |

**Fig S29. Natural selection analysis on chondrichthyan GNGT1 genes.** PAML was used to compute the dN/dS (= ω) per branch under four different models. **A**, Free-ratio model which fits one ω ratio per branch. **B**, Two-ratio model with one ω ratio for *Somniosus microcephalus* and one ω ratio for other shark species. **C**, One-ratio model with the same ω ratio for all shark species. **D**, RELAX ω distribution results under the RELAX alternative model, assigning *Somniosus microcephalus* as test branches and all other shark species as reference branches. **E**, Log-likelihood (lnL) and number of parameters (np) of each model. **F**, Log-likelihood ratio test (LRT) between each of the models, showing degrees of freedom (df) and statistical significance (p.value) for each comparison.

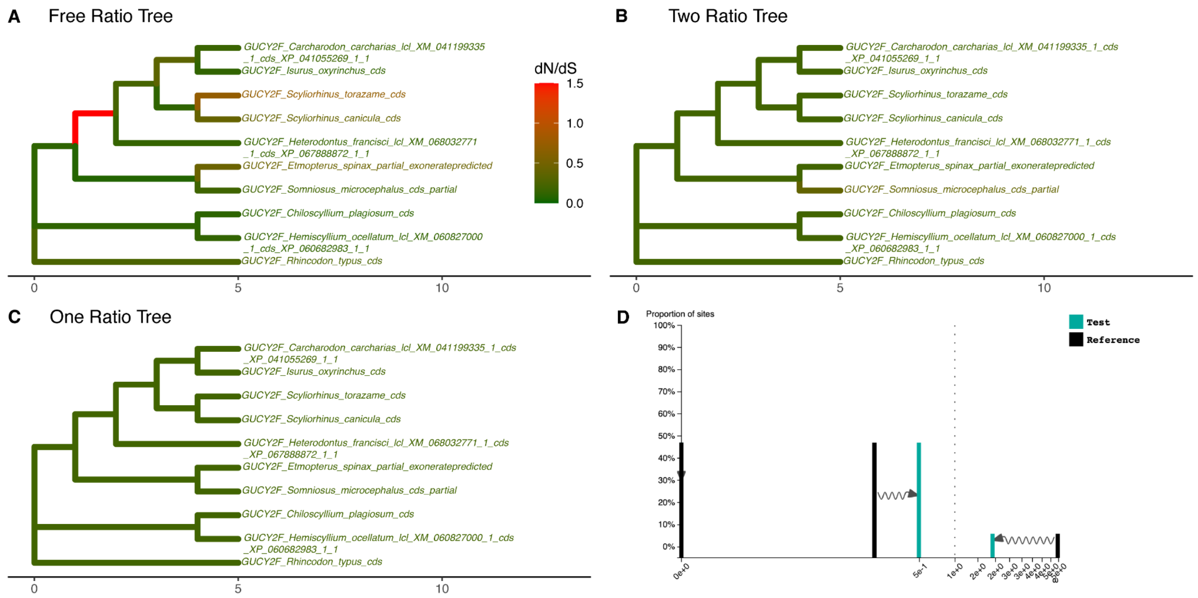


E

| Model | np | lnL |
|---|---|---|
| FreeRatio | 35 | −13999.66 |
| OneRatio | 19 | −14093.20 |
| TwoRatio | 20 | −14092.10 |

F

| Comparison | LRT | df | p.value |
|---|---|---|---|
| TwoRatio vs OneRatio | 2.19824 | 1 | 1.381684e−01 |
| FreeRatio vs OneRatio | 187.07788 | 16 | 3.194884e−31 |
| FreeRatio vs TwoRatio | 184.87964 | 15 | 2.461153e−31 |

**Fig S30. Natural selection analysis on chondrichthyan GUCY2F genes.** PAML was used to compute the dN/dS (= ω) per branch under four different models. **A**, Free-ratio model which fits one ω ratio per branch. **B**, Two-ratio model with one ω ratio for *Somniosus microcephalus* and one ω ratio for other shark species. **C**, One-ratio model with the same ω ratio for all shark species. **D**, RELAX ω distribution results under the RELAX alternative model, assigning *Somniosus microcephalus* as test branches and all other shark species as reference branches. **E**, Log-likelihood (lnL) and number of parameters (np) of each model. **F**, Log-likelihood ratio test (LRT) between each of the models, showing degrees of freedom (df) and statistical significance (p.value) for each comparison.

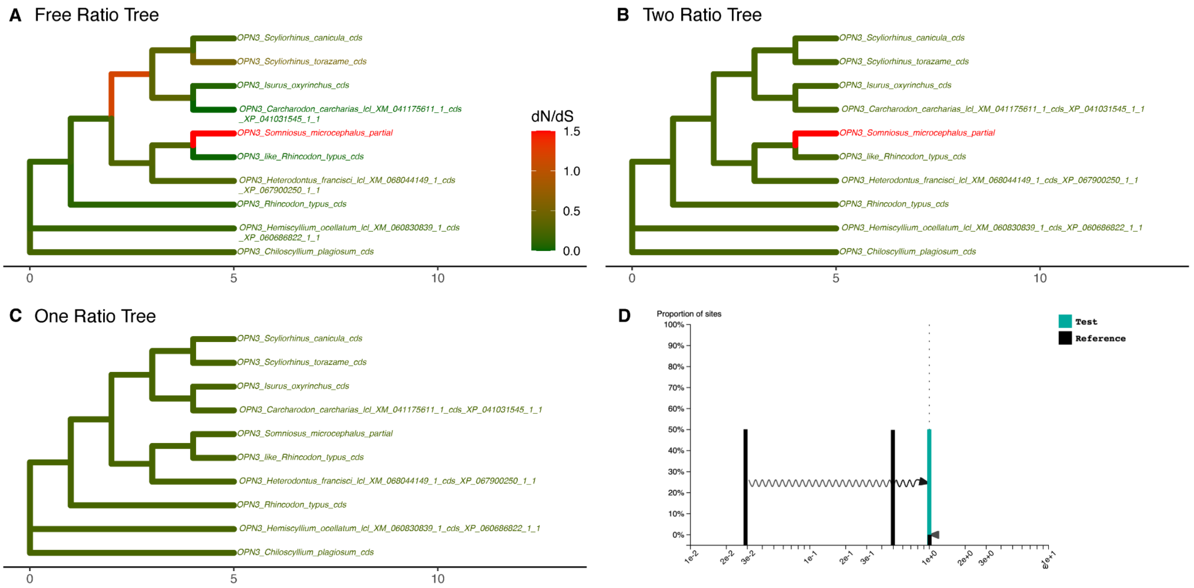


E

| Model | np | lnL |
|---|---|---|
| FreeRatio | 35 | −5166.571 |
| OneRatio | 19 | −5189.346 |
| TwoRatio | 20 | −5188.672 |

F

| Comparison | LRT | df | p.value |
|---|---|---|---|
| TwoRatio vs OneRatio | 1.348642 | 1 | 0.2455156645 |
| FreeRatio vs OneRatio | 45.550090 | 16 | 0.0001142421 |
| FreeRatio vs TwoRatio | 44.201448 | 15 | 0.0001022561 |

**Fig S31. Natural selection analysis on chondrichthyan OPN3 genes.** PAML was used to compute the dN/dS (= ω) per branch under four different models. **A**, Free-ratio model which fits one ω ratio per branch. **B**, Two-ratio model with one ω ratio for *Somniosus microcephalus* and one ω ratio for other shark species. **C**, One-ratio model with the same ω ratio for all shark species. **D**, RELAX ω distribution results under the RELAX alternative model, assigning *Somniosus microcephalus* as test branches and all other shark species as reference branches. **E**, Log-likelihood (lnL) and number of parameters (np) of each model. **F**, Log-likelihood ratio test (LRT) between each of the models, showing degrees of freedom (df) and statistical significance (p.value) for each comparison.

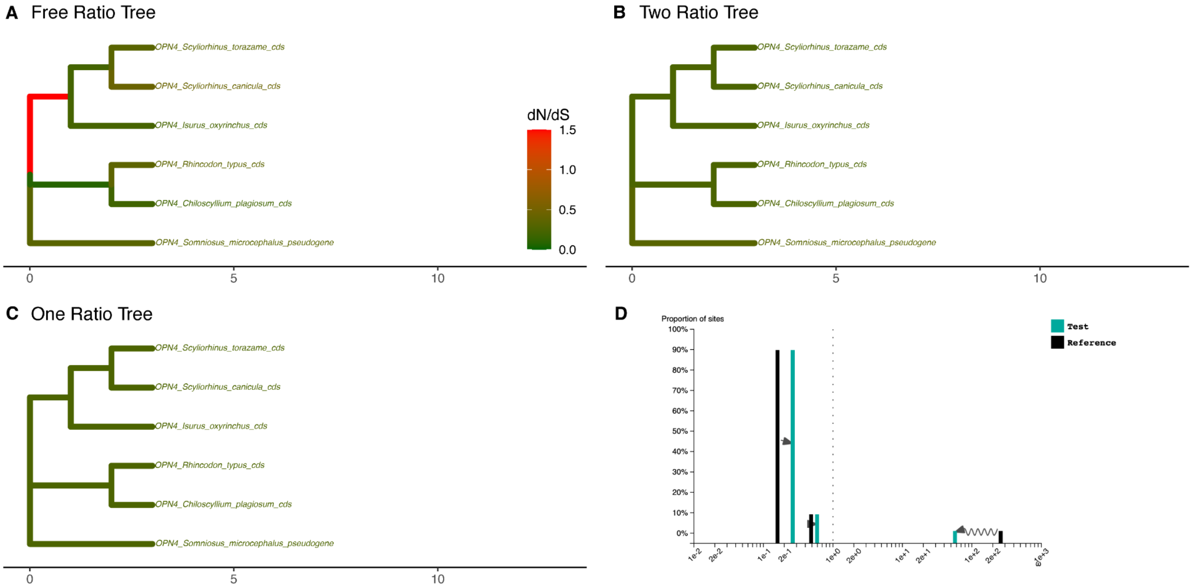


E

| Model | np | lnL |
|---|---|---|
| FreeRatio | 19 | –4275.333 |
| OneRatio | 11 | –4283.695 |
| TwoRatio | 12 | –4283.389 |

F

| Comparison | LRT | df | p.value |
|---|---|---|---|
| TwoRatio vs OneRatio | 0.611948 | 1 | 0.43405530 |
| FreeRatio vs OneRatio | 16.724882 | 8 | 0.03310482 |
| FreeRatio vs TwoRatio | 16.112934 | 7 | 0.02410437 |

**Fig S32. Natural selection analysis on chondrichthyan OPN4 genes.** PAML was used to compute the dN/dS (= ω) per branch under four different models. **A**, Free-ratio model which fits one ω ratio per branch. **B**, Two-ratio model with one ω ratio for *Somniosus microcephalus* and one ω ratio for other shark species. **C**, One-ratio model with the same ω ratio for all shark species. **D**, RELAX ω distribution results under the RELAX alternative model, assigning *Somniosus microcephalus* as test branches and all other shark species as reference branches. **E**, Log-likelihood (lnL) and number of parameters (np) of each model. **F**, Log-likelihood ratio test (LRT) between each of the models, showing degrees of freedom (df) and statistical significance (p.value) for each comparison.

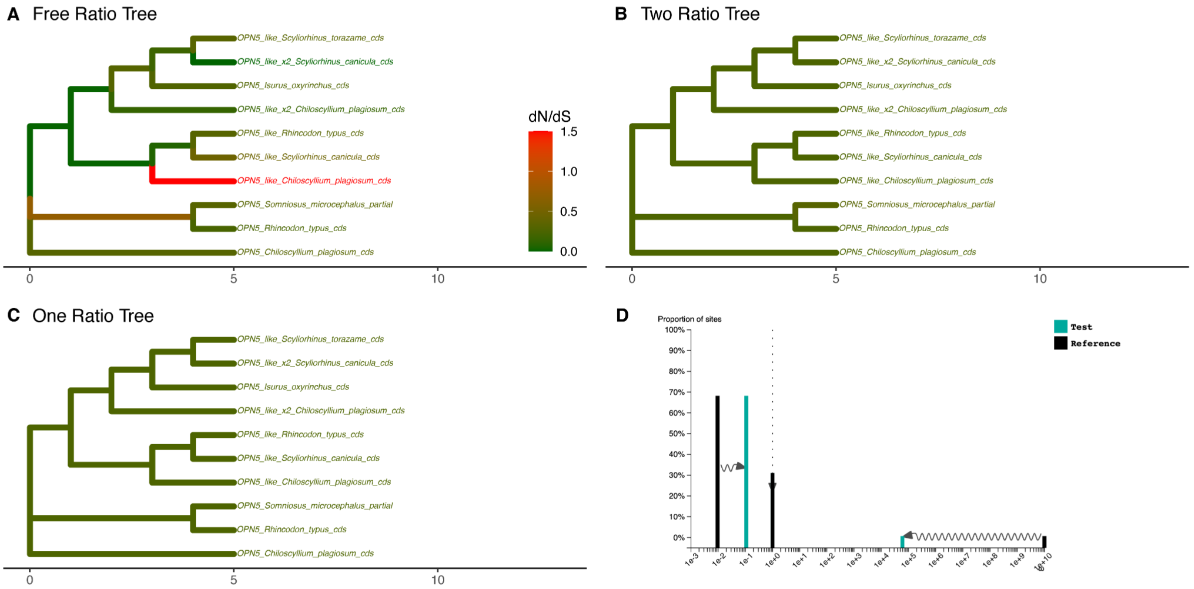


E

| Model | np | lnL |
|---|---|---|
| FreeRatio | 35 | −7271.134 |
| OneRatio | 19 | −7303.210 |
| TwoRatio | 20 | −7302.802 |

F

| Comparison | LRT | df | p.value |
|---|---|---|---|
| TwoRatio vs OneRatio | 0.81711 | 1 | 3.660265e−01 |
| FreeRatio vs OneRatio | 64.15175 | 16 | 1.030329e−07 |
| FreeRatio vs TwoRatio | 63.33464 | 15 | 6.677563e−08 |

**Fig S33. Natural selection analysis on chondrichthyan OPN5 genes.** PAML was used to compute the dN/dS (= ω) per branch under four different models. **A**, Free-ratio model which fits one ω ratio per branch. **B**, Two-ratio model with one ω ratio for *Somniosus microcephalus* and one ω ratio for other shark species. **C**, One-ratio model with the same ω ratio for all shark species. **D**, RELAX ω distribution results under the RELAX alternative model, assigning *Somniosus microcephalus* as test branches and all other shark species as reference branches. **E**, Log-likelihood (lnL) and number of parameters (np) of each model. **F**, Log-likelihood ratio test (LRT) between each of the models, showing degrees of freedom (df) and statistical significance (p.value) for each comparison.

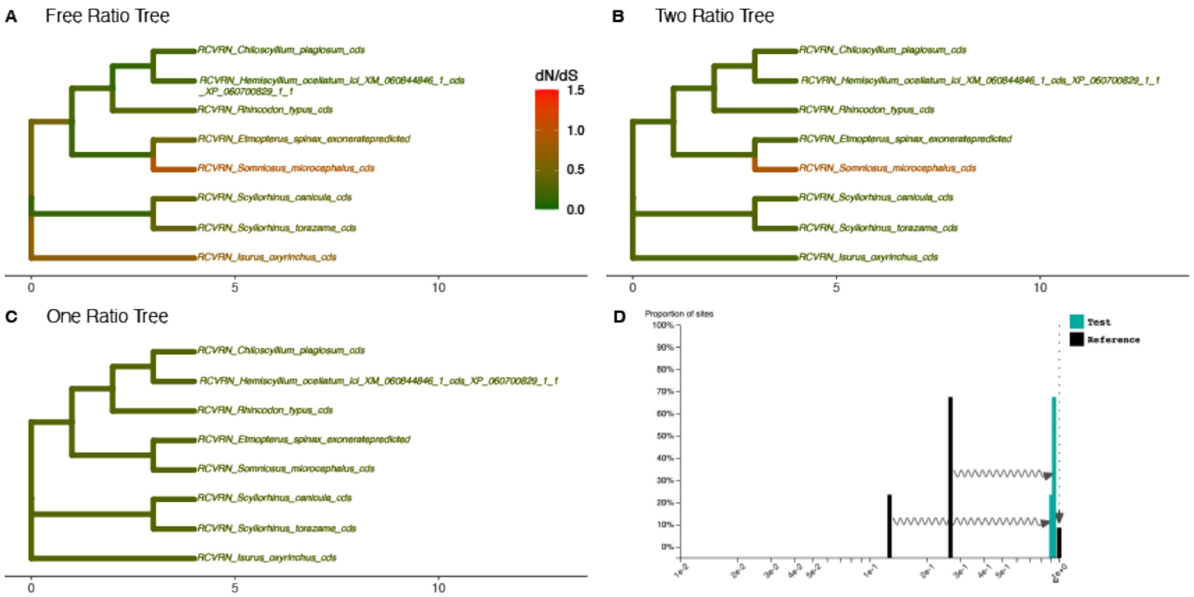


E

| Model | np | lnL |
|---|---|---|
| FreeRatio | 27 | −1841.589 |
| OneRatio | 15 | −1849.292 |
| TwoRatio | 16 | −1847.410 |

F

| Comparison | LRT | df | p.value |
|---|---|---|---|
| TwoRatio vs OneRatio | 3.76332 | 1 | 0.05238846 |
| FreeRatio vs OneRatio | 15.40496 | 12 | 0.22003392 |
| FreeRatio vs TwoRatio | 11.64164 | 11 | 0.39117389 |

**Fig S34. Natural selection analysis on chondrichthyan RCVRN genes.** PAML was used to compute the dN/dS (= ω) per branch under four different models. **A**, Free-ratio model which fits one ω ratio per branch. **B**, Two-ratio model with one ω ratio for *Somniosus microcephalus* and one ω ratio for other shark species. **C**, One-ratio model with the same ω ratio for all shark species. **D**, RELAX ω distribution results under the RELAX alternative model, assigning *Somniosus microcephalus* as test branches and all other shark species as reference branches. **E**, Log-likelihood (lnL) and number of parameters (np) of each model. **F**, Log-likelihood ratio test (LRT) between each of the models, showing degrees of freedom (df) and statistical significance (p.value) for each comparison.

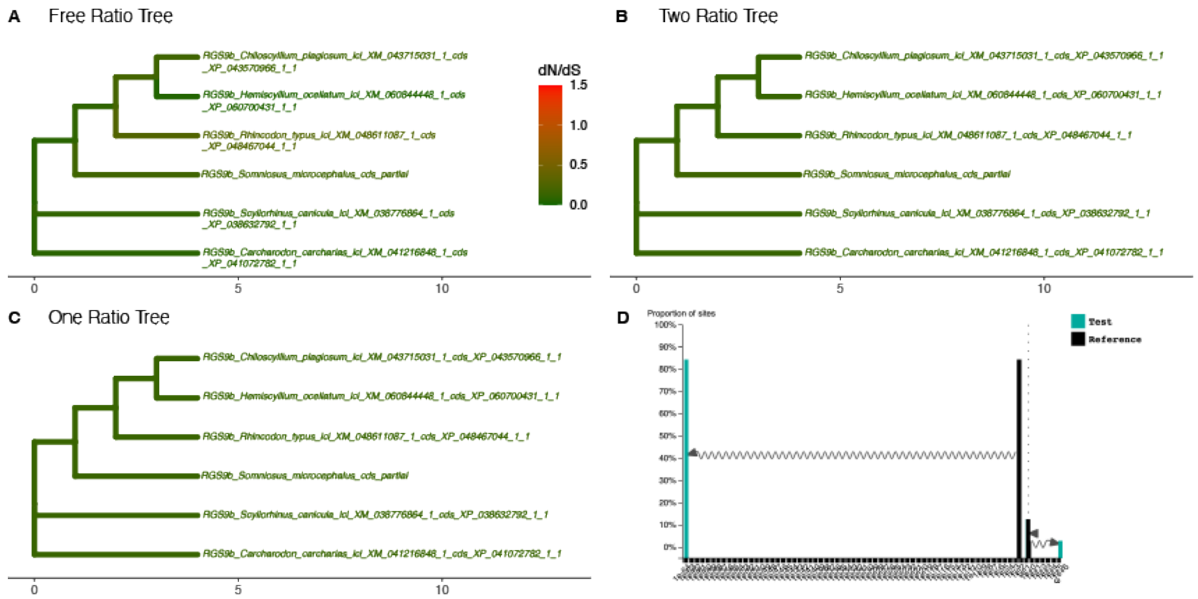


E

| Model | np | lnL |
|---|---|---|
| FreeRatio | 19 | −5532.320 |
| OneRatio | 11 | −5541.759 |
| TwoRatio | 12 | −5541.333 |

F

| Comparison | LRT | df | p.value |
|---|---|---|---|
| TwoRatio vs OneRatio | 0.851874 | 1 | 0.35602273 |
| FreeRatio vs OneRatio | 18.877902 | 8 | 0.01552644 |
| FreeRatio vs TwoRatio | 18.026028 | 7 | 0.01185312 |

**Fig S35. Natural selection analysis on chondrichthyan RGS9b genes.** PAML was used to compute the dN/dS (= ω) per branch under four different models. **A**, Free-ratio model which fits one ω ratio per branch. **B**, Two-ratio model with one ω ratio for *Somniosus microcephalus* and one ω ratio for other shark species. **C**, One-ratio model with the same ω ratio for all shark species. **D**, RELAX ω distribution results under the RELAX alternative model, assigning *Somniosus microcephalus* as test branches and all other shark species as reference branches. **E**, Log-likelihood (lnL) and number of parameters (np) of each model. **F**, Log-likelihood ratio test (LRT) between each of the models, showing degrees of freedom (df) and statistical significance (p.value) for each comparison.

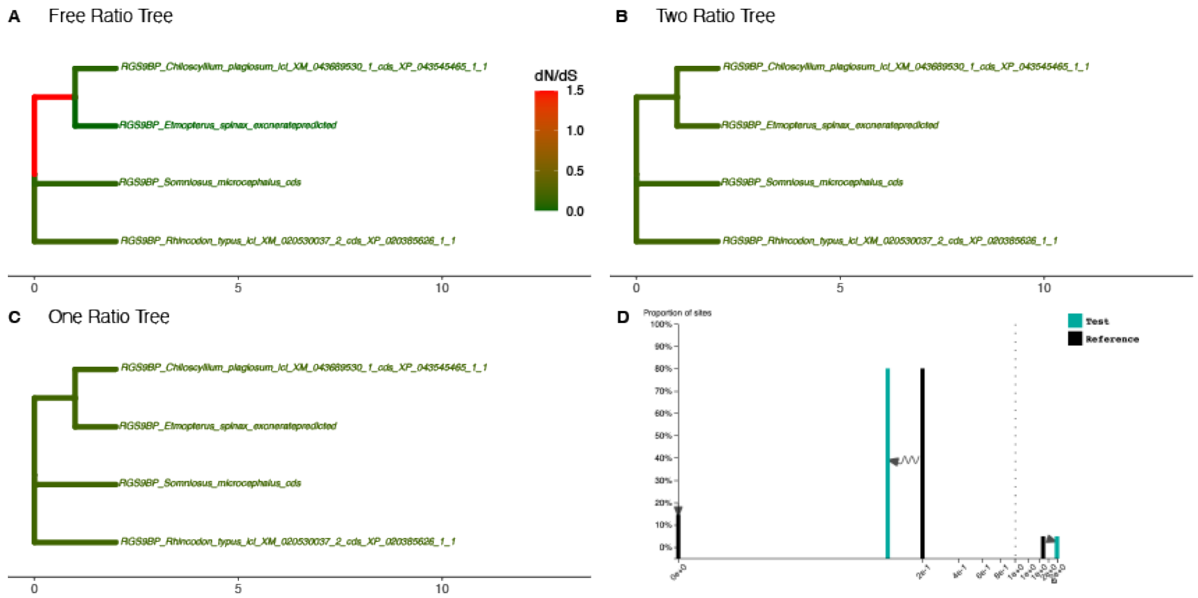


E

| Model | np | lnL |
|---|---|---|
| FreeRatio | 11 | −2237.273 |
| OneRatio | 7 | −2240.903 |
| TwoRatio | 8 | −2240.879 |

F

| Comparison | LRT | df | p.value |
|---|---|---|---|
| TwoRatio vs OneRatio | 0.048556 | 1 | 0.82559526 |
| FreeRatio vs OneRatio | 7.259642 | 4 | 0.12278716 |
| FreeRatio vs TwoRatio | 7.211086 | 3 | 0.06546557 |

**Fig S36. Natural selection analysis on chondrichthyan RGS9BP genes.** PAML was used to compute the dN/dS (= ω) per branch under four different models. **A**, Free-ratio model which fits one ω ratio per branch. **B**, Two-ratio model with one ω ratio for *Somniosus microcephalus* and one ω ratio for other shark species. **C**, One-ratio model with the same ω ratio for all shark species. **D**, RELAX ω distribution results under the RELAX alternative model, assigning *Somniosus microcephalus* as test branches and all other shark species as reference branches. **E**, Log-likelihood (lnL) and number of parameters (np) of each model. **F**, Log-likelihood ratio test (LRT) between each of the models, showing degrees of freedom (df) and statistical significance (p.value) for each comparison.

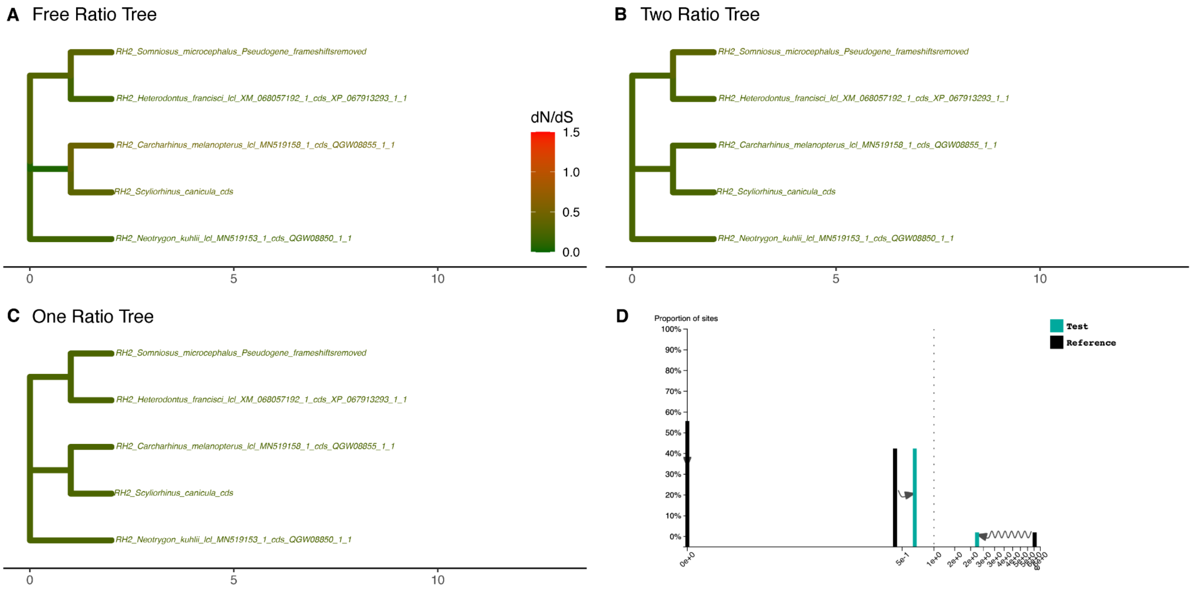


E

| Model | np | lnL |
|---|---|---|
| FreeRatio | 15 | –3053.678 |
| OneRatio | 9 | –3061.779 |
| TwoRatio | 10 | –3061.342 |

F

| Comparison | LRT | df | p.value |
|---|---|---|---|
| TwoRatio vs OneRatio | 0.873332 | 1 | 0.350034519 |
| FreeRatio vs OneRatio | 16.202466 | 6 | 0.012707536 |
| FreeRatio vs TwoRatio | 15.329134 | 5 | 0.009044794 |

**Fig S37. Natural selection analysis on chondrichthyan RH2 genes.** PAML was used to compute the dN/dS (= ω) per branch under four different models. **A**, Free-ratio model which fits one ω ratio per branch. **B**, Two-ratio model with one ω ratio for *Somniosus microcephalus* and one ω ratio for other shark species. **C**, One-ratio model with the same ω ratio for all shark species. **D**, RELAX ω distribution results under the RELAX alternative model, assigning *Somniosus microcephalus* as test branches and all other shark species as reference branches. **E**, Log-likelihood (lnL) and number of parameters (np) of each model. **F**, Log-likelihood ratio test (LRT) between each of the models, showing degrees of freedom (df) and statistical significance (p.value) for each comparison.

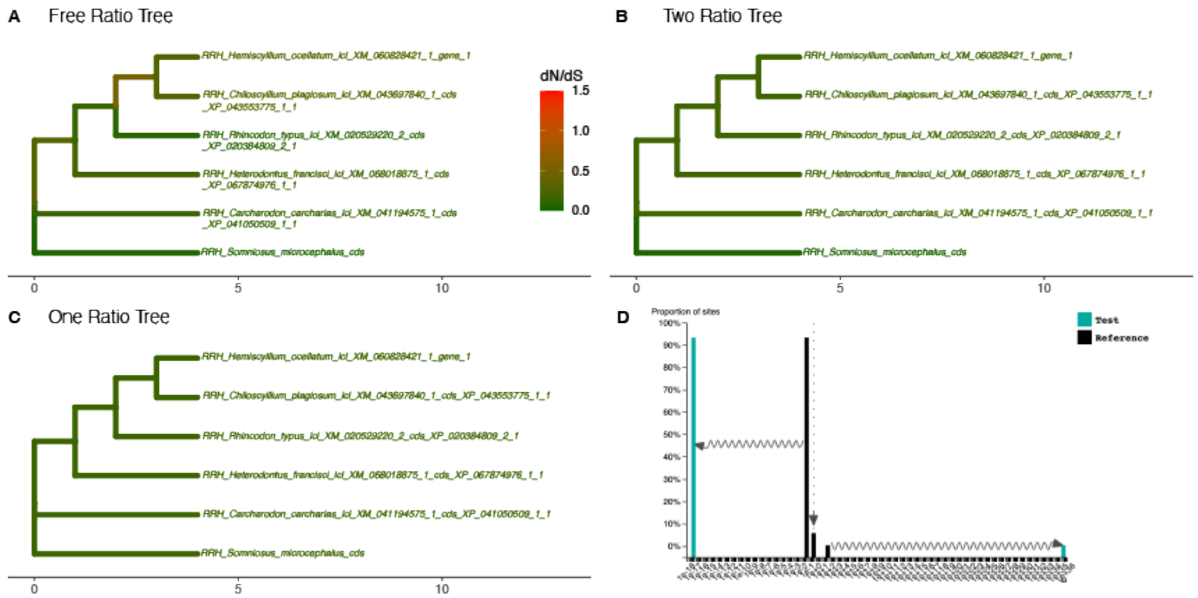


E

| Model | np | lnL |
|---|---|---|
| FreeRatio | 19 | −2377.464 |
| OneRatio | 11 | −2385.849 |
| TwoRatio | 12 | −2382.763 |

F

| Comparison | LRT | df | p.value |
|---|---|---|---|
| TwoRatio vs OneRatio | 6.171354 | 1 | 0.01298352 |
| FreeRatio vs OneRatio | 16.768736 | 8 | 0.03260935 |
| FreeRatio vs TwoRatio | 10.597382 | 7 | 0.15717066 |

**Fig S38. Natural selection analysis on chondrichthyan RRH genes.** PAML was used to compute the dN/dS (= ω) per branch under four different models. **A**, Free-ratio model which fits one ω ratio per branch. **B**, Two-ratio model with one ω ratio for *Somniosus microcephalus* and one ω ratio for other shark species. **C**, One-ratio model with the same ω ratio for all shark species. **D**, RELAX ω distribution results under the RELAX alternative model, assigning *Somniosus microcephalus* as test branches and all other shark species as reference branches. **E**, Log-likelihood (lnL) and number of parameters (np) of each model. **F**, Log-likelihood ratio test (LRT) between each of the models, showing degrees of freedom (df) and statistical significance (p.value) for each comparison.

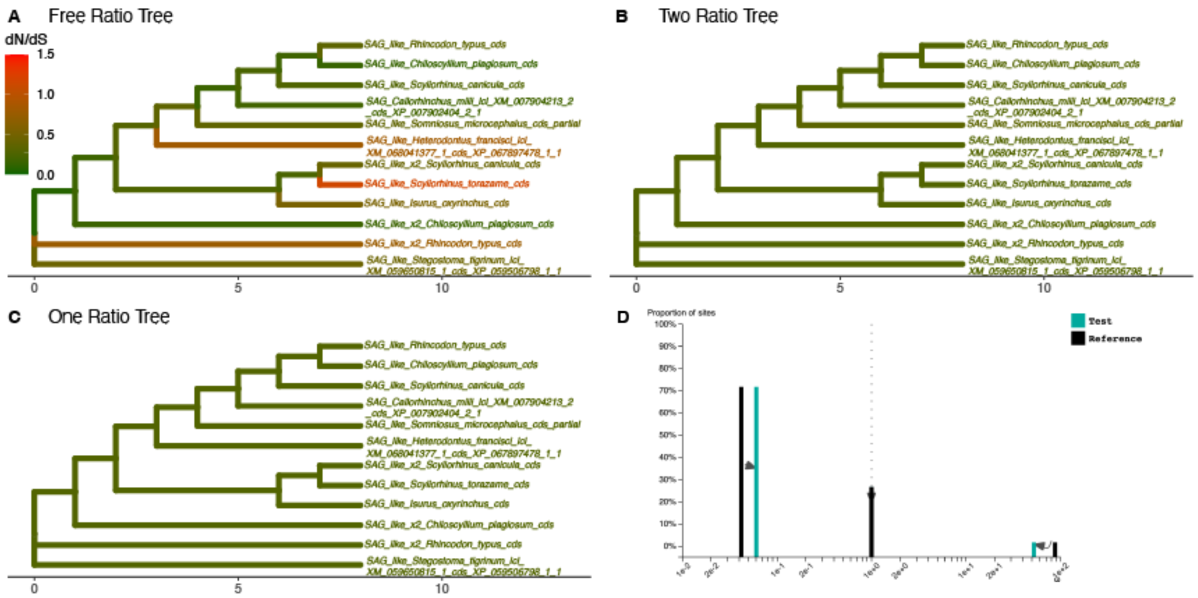


E

| Model | np | lnL |
|---|---|---|
| FreeRatio | 43 | −7273.097 |
| OneRatio | 23 | −7300.511 |
| TwoRatio | 24 | −7300.437 |

F

| Comparison | LRT | df | p.value |
|---|---|---|---|
| TwoRatio vs OneRatio | 0.14669 | 1 | 7.017188e−01 |
| FreeRatio vs OneRatio | 54.82663 | 20 | 4.358372e−05 |
| FreeRatio vs TwoRatio | 54.67994 | 19 | 2.601721e−05 |

**Fig S39. Natural selection analysis on chondrichthyan SAG genes.** PAML was used to compute the dN/dS (= ω) per branch under four different models. **A**, Free-ratio model which fits one ω ratio per branch. **B**, Two-ratio model with one ω ratio for *Somniosus microcephalus* and one ω ratio for other shark species. **C**, One-ratio model with the same ω ratio for all shark species. **D**, RELAX ω distribution results under the RELAX alternative model, assigning *Somniosus microcephalus* as test branches and all other shark species as reference branches. **E**, Log-likelihood (lnL) and number of parameters (np) of each model. **F**, Log-likelihood ratio test (LRT) between each of the models, showing degrees of freedom (df) and statistical significance (p.value) for each comparison.

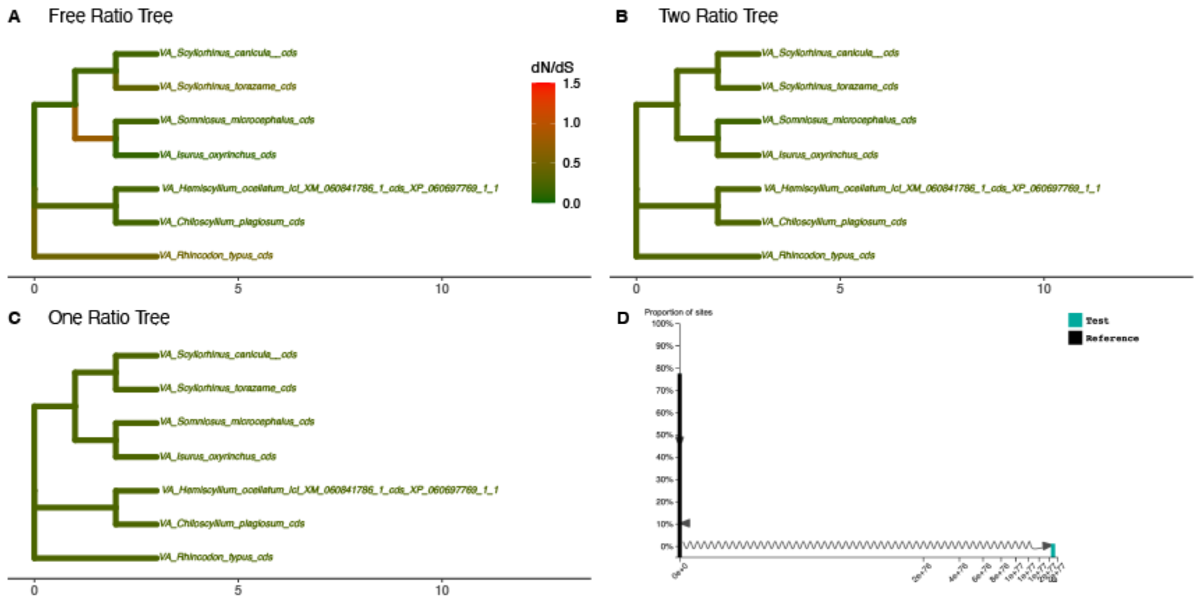


E

| Model | np | lnL |
|---|---|---|
| FreeRatio | 23 | −3777.448 |
| OneRatio | 13 | −3787.026 |
| TwoRatio | 14 | −3786.449 |

F

| Comparison | LRT | df | p.value |
|---|---|---|---|
| TwoRatio vs OneRatio | 1.153394 | 1 | 0.28283979 |
| FreeRatio vs OneRatio | 19.156192 | 10 | 0.03832307 |
| FreeRatio vs TwoRatio | 18.002798 | 9 | 0.03514109 |

**Fig S40. Natural selection analysis on chondrichthyan VA genes.** PAML was used to compute the dN/dS (= ω) per branch under four different models. **A**, Free-ratio model which fits one ω ratio per branch. **B**, Two-ratio model with one ω ratio for *Somniosus microcephalus* and one ω ratio for other shark species. **C**, One-ratio model with the same ω ratio for all shark species. **D**, RELAX ω distribution results under the RELAX alternative model, assigning *Somniosus microcephalus* as test branches and all other shark species as reference branches. **E**, Log-likelihood (lnL) and number of parameters (np) of each model. **F**, Log-likelihood ratio test (LRT) between each of the models, showing degrees of freedom (df) and statistical significance (p.value) for each comparison.

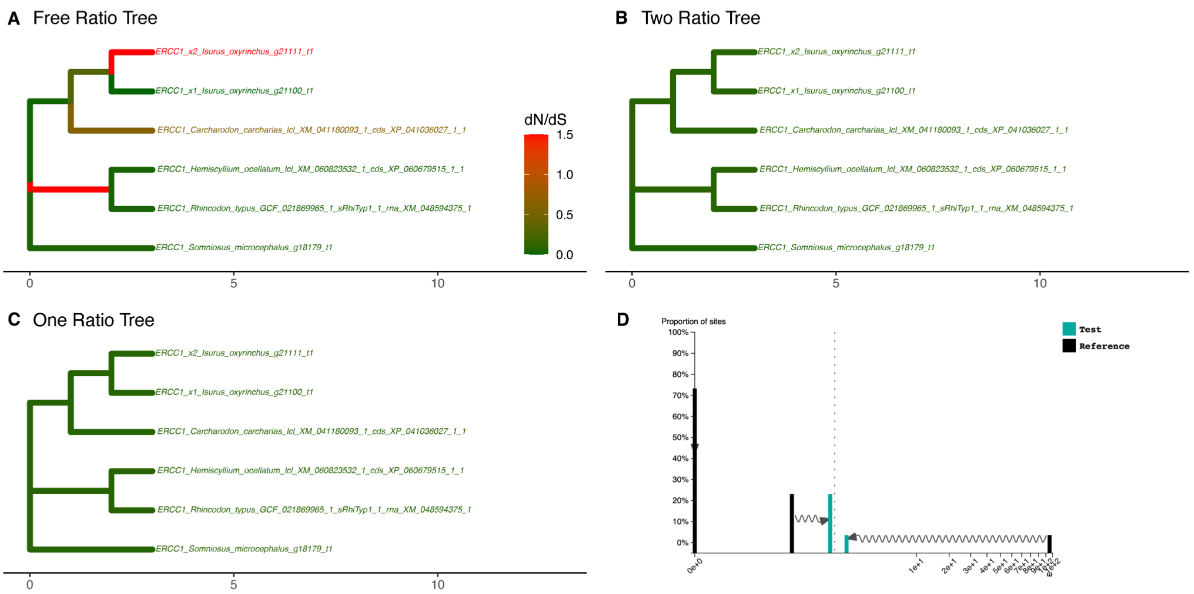


| E | Model | np | lnL |
|---|---|---|---|
| | FreeRatio | 19 | −1995.449 |
| | OneRatio | 11 | −2013.777 |
| | TwoRatio | 12 | −2012.890 |

| F | Comparison | LRT | df | p.value |
|---|---|---|---|---|
| | TwoRatio vs OneRatio | 1.77405 | 1 | 1.828817e−01 |
| | FreeRatio vs OneRatio | 36.65610 | 8 | 1.331175e−05 |
| | FreeRatio vs TwoRatio | 34.88205 | 7 | 1.176998e−05 |

**Fig S41. Natural selection analysis on chondrichthyan ERCC1 genes.** PAML was used to compute the dN/dS (= ω) per branch under four different models. **A**, Free-ratio model which fits one ω ratio per branch. **B**, Two-ratio model with one ω ratio for *Somniosus microcephalus* and one ω ratio for other shark species. **C**, One-ratio model with the same ω ratio for all shark species. **D**, RELAX ω distribution results under the RELAX alternative model, assigning *Somniosus microcephalus* as test branches and all other shark species as reference branches. **E**, Log-likelihood (lnL) and number of parameters (np) of each model. **F**, Log-likelihood ratio test (LRT) between each of the models, showing degrees of freedom (df) and statistical significance (p.value) for each comparison.

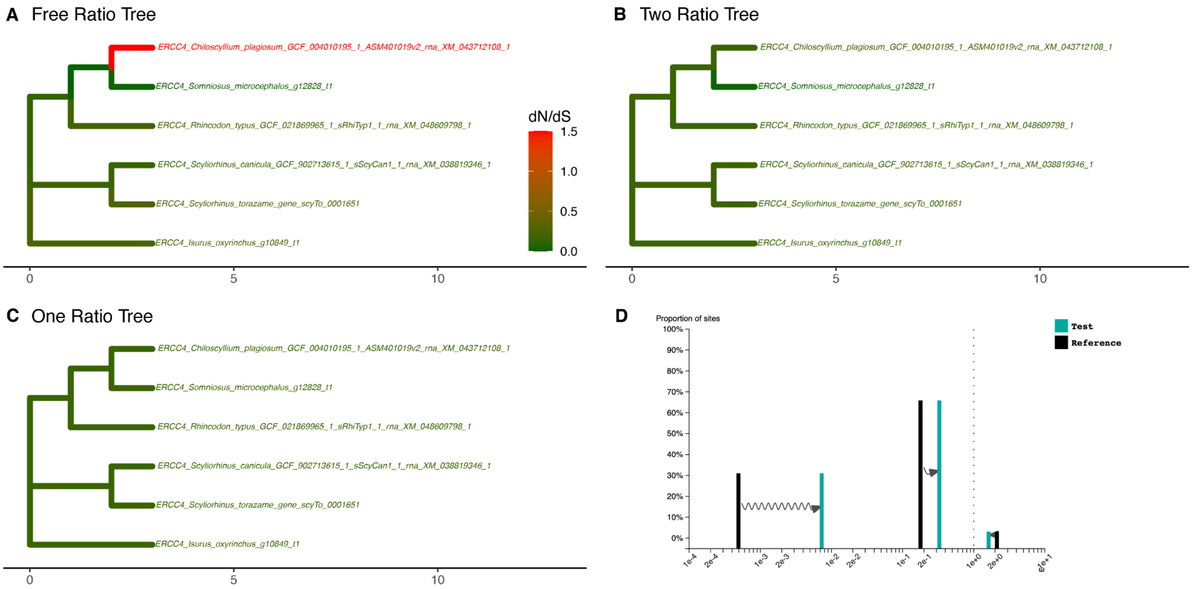


| E | | |
|---|---|---|
| Model | np | lnL |
| FreeRatio | 19 | −7299.413 |
| OneRatio | 11 | −7303.924 |
| TwoRatio | 12 | −7302.955 |

| F | | | |
|---|---|---|---|
| Comparison | LRT | df | p.value |
| TwoRatio vs OneRatio | 1.937958 | 1 | 0.1638904 |
| FreeRatio vs OneRatio | 9.021280 | 8 | 0.3405040 |
| FreeRatio vs TwoRatio | 7.083322 | 7 | 0.4202563 |

**Fig S42. Natural selection analysis on chondrichthyan ERCC4 genes.** PAML was used to compute the dN/dS (= ω) per branch under four different models. **A**, Free-ratio model which fits one ω ratio per branch. **B**, Two-ratio model with one ω ratio for *Somniosus microcephalus* and one ω ratio for other shark species. **C**, One-ratio model with the same ω ratio for all shark species. **D**, RELAX ω distribution results under the RELAX alternative model, assigning *Somniosus microcephalus* as test branches and all other shark species as reference branches. **E**, Log-likelihood (lnL) and number of parameters (np) of each model. **F**, Log-likelihood ratio test (LRT) between each of the models, showing degrees of freedom (df) and statistical significance (p.value) for each comparison.

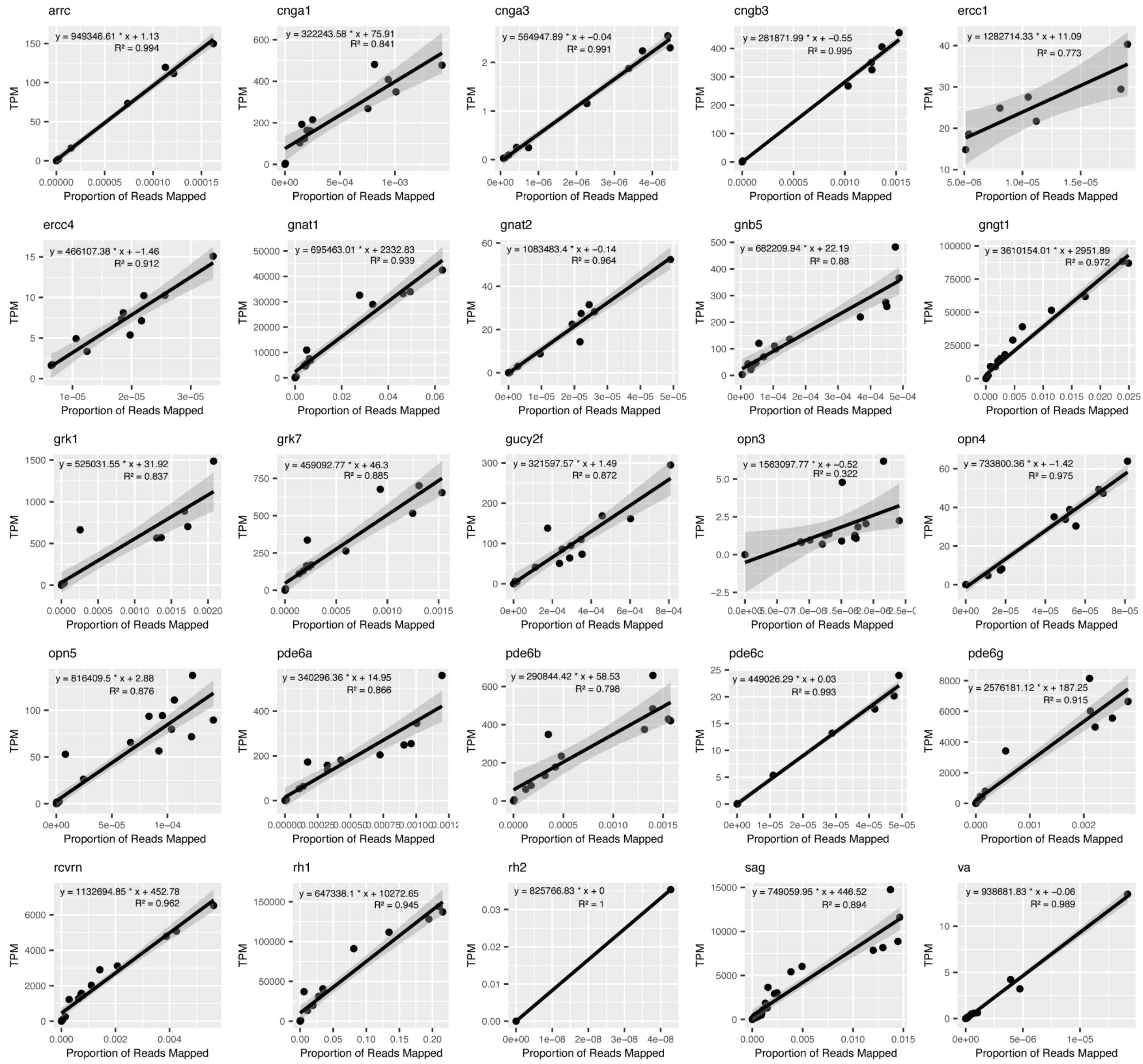


**Fig S43. Per-gene correlations between measures of gene expression.** For each gene analysed in this study, transcripts per-million (TPM) is plotted against the proportion of reads mapped to each gene in up to five shark species (*Scyliorhinus canicula, S. torazame, Isurus oxyrinchus, Rhincodon typus,* and *Chiloscyllium plagiosum*). The per-gene equations describing the linear regressions were used to extrapolate the TPM values for *Somniosus microcephalus* from the proportion of reads mapped.

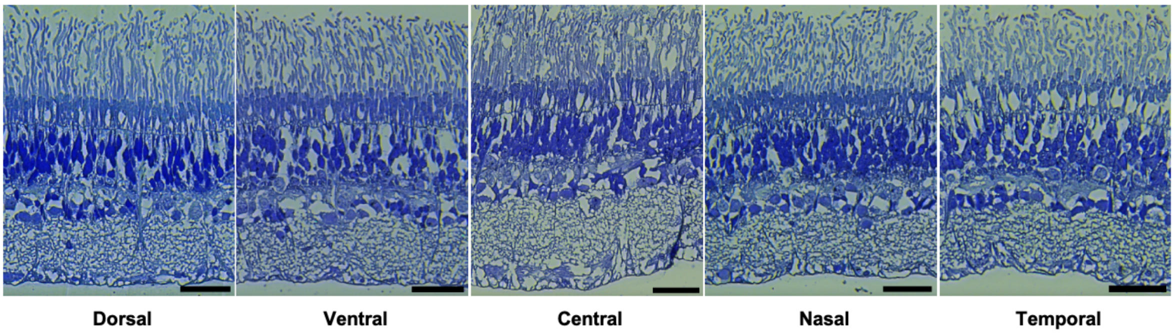


**Fig S44. Retinal architecture in different regions of the Greenland shark retina.** Transverse sections through five regions of the retina of *Somniosus microcephalus.* Scale bars: 50 µm.

**Table S1. Details of specimens used in this study.** All individuals were adult Greenland sharks (*Somniosus microcephalus*) collected off the coast of Greenland. M, male; F, female; U, unknown.

| **Sex** | **TL (m)** | **Specimen ID** | **Analyses** |
|---|---|---|---|
| M | 3.35 | GS24_6 | Transcriptome |
| F | 3.65 | GS24_9 | Transcriptome |
| F | 3.6 | GS24_12 | Transcriptome |
| M | 3.14 | GS23_1 | Ultramicrotomy |
| F | 3.09 | GS23_6 | Ultramicrotomy |
| F | 3.54 | GS24_4 | Ultramicrotomy |
| F | 3.31 | GS23_6 | RNAscope |
| U | U | Shark3 | Genome |

**Table S2. Genomes and transcriptomes used in this study.** All genomes are available from the NCBI Genome database and all transcriptomes are available from the Sequence Read Archive.

| **Species** | **Data type** | **Accession no.** |
|---|---|---|
| *Somniosus microcephalus* | Genome | SRR32965274 |
| | Transcriptome - retina | SRR32965277 |
| | Transcriptome - retina | SRR32965276 |
| | Transcriptome - retina | SRR32965275 |
| *Scyliorhinus torazame* | Genome | GCA_003427355.1 |
| | Transcriptome - retina | DRR111795 |
| | Transcriptome - retina | DRR111817 |
| *Scyliorhinus canicula* | Genome | GCF_902713615.1 |
| | Transcriptome - retina | SRR12813952 |
| | Transcriptome - retina | SRR12813953 |
| | Transcriptome - retina | SRR12813954 |
| | Transcriptome - retina | SRR12813957 |
| | Transcriptome - retina | SRR12813958 |
| *Rhincodon typus* | Genome | GCF_021869965.1 |
| | Transcriptome - retina | SRR19140294 |
| *Isurus oxyrinchus* | Genome | GCA_026770705.1 |
| | Transcriptome - retina | SRR10998462 |
| | Transcriptome - retina | SRR10998463 |

| | | |
|---|---|---|
| | Transcriptome - retina | SRR10998464 |
| *Chiloscyllium plagiosum* | Genome | GCF_004010195.1 |
| | Transcriptome - retina | SRR7539029 |
| | Transcriptome - retina | SRR7539030 |
| | Transcriptome - retina | SRR7539034 |
| | Transcriptome - retina | SRR7539035 |
| | Transcriptome - retina | SRR7539036 |
| *Amblyraja radiata* | Genome | GCA_010909765.2 |
| *Callorhinchus milii* | Genome | GCA_018977255.1 |
| *Carcharodon carcharias* | Genome | GCA_017639515.1 |
| *Chiloscyllium punctatum* | Genome | GCA_003427335.1 |
| *Ginglymostoma cirratum* | Genome | GCA_024137785.1 |
| *Hemiscyllium ocellatum* | Genome | GCA_020745735.1 |
| *Leucoraja erinacea* | Genome | GCA_028641065.1 |
| *Pristis pectinata* | Genome | GCA_009764475.2 |
| *Raja brachyura* | Genome | GCA_963514005.1 |
| *Sphyrna mokarran* | Genome | GCA_024679065.1 |
| *Squalus acanthias* | Genome | GCA_030390025.1 |

**Table S3. Assembly statistics for the Greenland shark genome.** Statistics generated for a scaffold-level assembly using the *de novo* sequence assembler, ABySS.

| | |
|---|---|
| **Total number of scaffolds (n)** | 29940000 |
| **Number of sequences that are at least 500 bp long (n:500)** | 1175830 |
| **L50** | 191587 |
| **Length of shortest sequence (min)** | 500 |
| **N75** | 1623 |
| **N50** | 3383 |
| **N25** | 6343 |
| **Effective size (E-size)** | 4810 |
| **Length of longest sequence (max)** | 149844 |
| **Total length of all sequences in the assembly (sum)** | 2365000000 |
| **BUSCO complete (%)** | 14.2 |
| **BUSCO fragmented (%)** | 41.0 |
| **BUSCO missing (%)** | 44.8 |

**Table S4. RNAscope probes.** Details of probes used for RNAscope experiments.

| RNAscope Probe | Catalog number |
|---|---|
| Smi-Rh1-C1 | 1595991-C1 |
| Smi-Gad1-C2 | 1596161-C2 |
| Smi-pkcalpha-C3 | 1595891-C3 |
| Smi-Glula-C1 | 1595881-C1 |
| Smi-RBPMS2-C2 | 1595851-C2 |
| Smi-ELOVL2-C3 | 1596121-C3 |
| Smi-Slc6a9-C | 1596131-C3 |
